# Carbon-to-ATP Ratios Across the Kingdoms of Life

**DOI:** 10.64898/2026.01.27.702009

**Authors:** Peyman Fahimi, Mohammad M. Amirian, Andrew J. Irwin, Zoe V. Finkel

## Abstract

ATP is the major energy-carrying molecule in cells, driving chemical reactions for cell development and growth. A diverse set of metabolically active marine bacteria and unicellular eukaryotes appear to maintain an approximately constant cellular carbon-to-ATP ratio (C:ATP) of 250 (g/g), which has been used as a predictor of marine microbial biomass for about 60 years. We have compiled ∼ 400 measurements of ATP and carbon content from more than 80 papers published between 1964 and 2024, spanning organisms from bacteria to animals and plants along with a wide range of tissues. These data show that the carbon-to-ATP ratio varies by a staggering six orders of magnitude, depending not only on physiological and environmental conditions but also on species, completely contradicting the long-standing assumption in the literature. Here we develop a theory for organismal C:ATP content based upon metabolic rate, ATP transportation time within the organism, and ratio of structural to functional carbon within an organism, accentuating why such a deviation from 250 should be expected. Our model predicts both the median and variation in C:ATP ratios across organisms across the tree of life. We find the median C:ATP is typically within a factor of two of 250 for bacteria and unicellular marine eukaryotes. The ratio is notably lower in multicellular animals and higher for leaves and roots of land plants. Eukaryotic photosynthetic organisms have a median C:ATP > 250, which we attribute mainly to the proximity of mitochondria and chloroplasts, both of which tend to be near ATP consumption sites, thereby reducing ATP transport time relative to heterotrophs. Within broad taxonomic groups we predict variation of six orders of magnitude in C:ATP due to differences in biomass-normalized metabolic rate and the amount of non-metabolically active, structural material in organisms.

## 1. Introduction

Understanding the energetic costs of building and operating living systems is essential to explaining patterns of growth, development, and ecological trade-offs across species [1,2]. These costs, measured in terms of ATP (Adenosine-5’-triphosphate) molecules that fuel most cellular processes [3], are fundamental to understand how organisms allocate resources and function. A central question is how much ATP mass is available per unit of metabolically active carbon, expressed as the C:ATP mass ratio. The long-standing benchmark of ∼250 for microbes has served as a proxy to estimate living biomass in marine systems, with deviations typically attributed to detritus, dead cells, or methodological differences [4–7]. However, we compiled 400 data points from numerous independent studies published over the past 60 years (see Supplementary Section I) showing that this ratio varies widely across species (highest in SAR11, lowest in the pupal brain of the cotton bollworm), physiological states (e.g., organism size, growth rate, carbon source, workload, stress exposure, developmental stage), and environmental conditions (e.g., nutrient and oxygen availability, temperature, salinity). We seek the physiological meaning behind this ratio and argue that such variability cannot be dismissed as experimental artifacts [5,8] but should be understood mechanistically.

Because our compiled data reveal a statistically significant relationship between C:ATP and organism or tissue size (see Supplementary Section I), we developed a size-dependent model of ATP and carbon content. Cellular ATP content 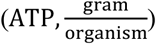 can be computed as the product of ATP consumption rate and the average time between ATP production and consumption, known as the turnover time (*τ, s*) [9]. Under steady-state conditions in actively growing cells, the ATP production rate 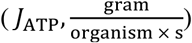 approximately matches the ATP consumption rate [9,10]. Therefore, ATP content can be expressed as the product of the ATP production rate and ATP turnover time (Eq. 1), linking ATP content to measurable parameters including respiration and photosynthesis rates [11–15], as well as biophysical constraints such as ATP transport time.

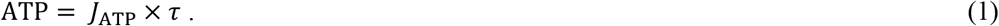

Although the links between respiration, photosynthesis, and ATP production rates are well stablished across taxa [16,17], and both respiration and photosynthesis scale with organism size [1,18], ATP turnover time remains poorly characterized, particularly in animals. Standard formulas for free diffusion and diffusive target search provide lower and upper bounds for size-dependent ATP turnover time, yet a more precise model would require detailed information on the copy number of mitochondria relative to cell volume, their spatial distribution within the 3D cellular space, the ATP consumption rate of different organelle types, and the packing density of cellular structures. These factors collectively determine how ATP is transported from mitochondria to sites of consumption, and thus the true ATP turnover time. Developing such a comprehensive model is beyond the scope of this paper. Instead, we address the ATP turnover time problem by combining empirical ATP content data with theoretical estimates of ATP production rates to infer turnover times, and then examine their biophysical underpinnings through cell size, mitochondrial positioning, and mitochondrial abundance. Carbon content is strongly size-dependent and is considerably more stable than ATP content, yet showing variation due to structural features such as cell walls and tissue type. By integrating these elements, we construct a mechanistic model that not only explains the median C:ATP ratio but also accounts for its variation within and across kingdoms as a universal metric.

## 2. Materials and Methods

ATP and carbon content can be modeled in heterotrophs and photoautotrophs, from bacteria to animals and plants, as the outcome of interactions among underlying biological, biochemical, and biophysical variables. Herein we design such a model and test it against an extensive database compiled from 60 years of literature across the kingdoms of life (see Supplementary Section II).

### 2.1. ATP Content

In aerobic heterotrophs, the majority of ATP (∼90%) is generated via oxidative phosphorylation, with the remainder produced through substrate-level phosphorylation in glycolysis and the small GTP yield of the tricarboxylic acid (TCA) cycle [16]. Within the mitochondrial matrix, NADH and FADH_2_ donate electrons to the electron transport chain, ultimately reducing oxygen to water. The released energy pumps protons across the membrane, and their return through ATP synthase drives ATP synthesis (with bacteria using an analogous chemiosmotic system operating across their cytoplasmic membrane) [3,19,20]. A portion of protons bypasses ATP synthase, re-entering the mitochondrial matrix without contributing to ATP synthesis, thereby dissipating energy as heat [21]. This phenomenon, known as proton leak (*PL*_1_), arises from various factors, including the high proton concentration in the intermembrane space, intrinsic properties of the phospholipid membrane, and presence of water wires and anion carrier proteins [22] (See the Supplementary Section III for factors affecting proton leak and the corresponding value of the leak). The stoichiometric ratio of oxygen consumption to ATP synthesis ( 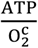; see Supplementary Section IV) [16] combined with the respiration rate 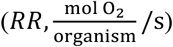 can be used to estimate the rate of ATP production. A proportion of respiration is lost due to proton leak, leaving *RR*(1 − *PL*_1_) as the effective respiration rate contributing to ATP production. A substantial body of literature demonstrates that respiration rate varies within species as a function of organism size [13,18] and growth rate [23–26], with further variation observed across species due to factors such as gene number, genome length, surface area limitations, and circulatory system differences [18]. We compiled data on heterotrophic and photoautotrophic organisms, from bacteria to mammals, to document the variation in respiration rate within and across species (See Supplementary Section II). Accordingly, the ATP production rate in heterotrophs can be expressed by the following equation, where the constant 507.2 represents the molar mass of ATP:

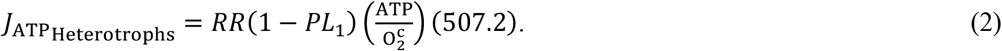

In photoautotrophs, additional processes (non-cyclic, cyclic and pseudo-cyclic electron transport, see Supplementary Section V) produce ATP [17,27]. Non-cyclic photophosphorylation in the chloroplast during photosynthesis involves a tightly regulated coupling between several key processes: the number of protons pumped into the thylakoid lumen, the number of oxygen molecules produced 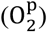, and the number of ATP molecules generated and required for carbon fixation [17,27]. The proportion of the total cellular ATP derived from photosynthesis versus respiration can be estimated using the dark respiration-to-photosynthesis ratio 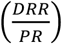 [28,29], which varies both within and across species (see Supplementary Section V). Protons leak across the thylakoid membrane (PL_2_), as also observed across the mitochondrial membrane. Assuming the dark respiration rate (*DRR*) of photoautotrophic organisms reported in Table S5 is equivalent to their light respiration rate, the total ATP production rate in these organisms can be determined as follows:

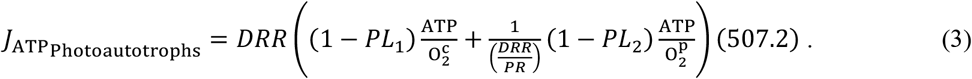

ATP turnover time (*τ* in Eq. 1) is the elapsed time from production to consumption of an ATP molecule averaged over the entire organism. If ATP is not readily available at the site of consumption, its turnover time will include the time required for transport from the site of production. Therefore, it is crucial to know the position of the mitochondrion and its distance from the sites of ATP consumption. ATP turnover time has been measured and reported in bacteria [9,30], microbial fungi [30], ciliates [31], and plants [30,32] while it remains less explored in animals. ATP turnover time can be measured experimentally; for example, using ^32^P radiolabeled phosphate tracer methods [33] or by tracking the ATP decline during the initial second of aminoethyldextrane (AED) stimulation [31].

While there are biological explanations for how ATP turnover time is regulated, including mechanisms maintaining the homeostatic balance among ATP, ADP, and AMP concentrations [34,35] there is no well-established physical mechanism that determines ATP turnover time. We address this problem using two complementary approaches. First, we compile data on ATP content across a wide range of organisms across the kingdoms of life (including ATP content in various animal tissues; Supplementary Section I) and divide it by our theoretically modeled ATP production rate, *J*_ATP_ (Eq. 1), to estimate ATP turnover time. We find that ATP turnover time is less than 1 second in microalgae, higher plants, and fast-growing bacteria, but it can vary by up to two orders of magnitude in heterotrophic protists and animals (Figure 1, Results section). To shed light on the biophysical basis of these estimated values, we use cell size data for unicellular organisms and the range of cell sizes in animals [36], along with literature reports on mitochondrial positioning [37,38] and population [39–41], and then examine how ATP diffusion time shapes overall ATP turnover time. To explore this further, we categorize organisms into three groups: (1) bacteria and cyanobacteria, (2) protists (both heterotrophic and photoautotrophic), and (3) multicellular organisms, including animals and plants.

**Figure 1:**
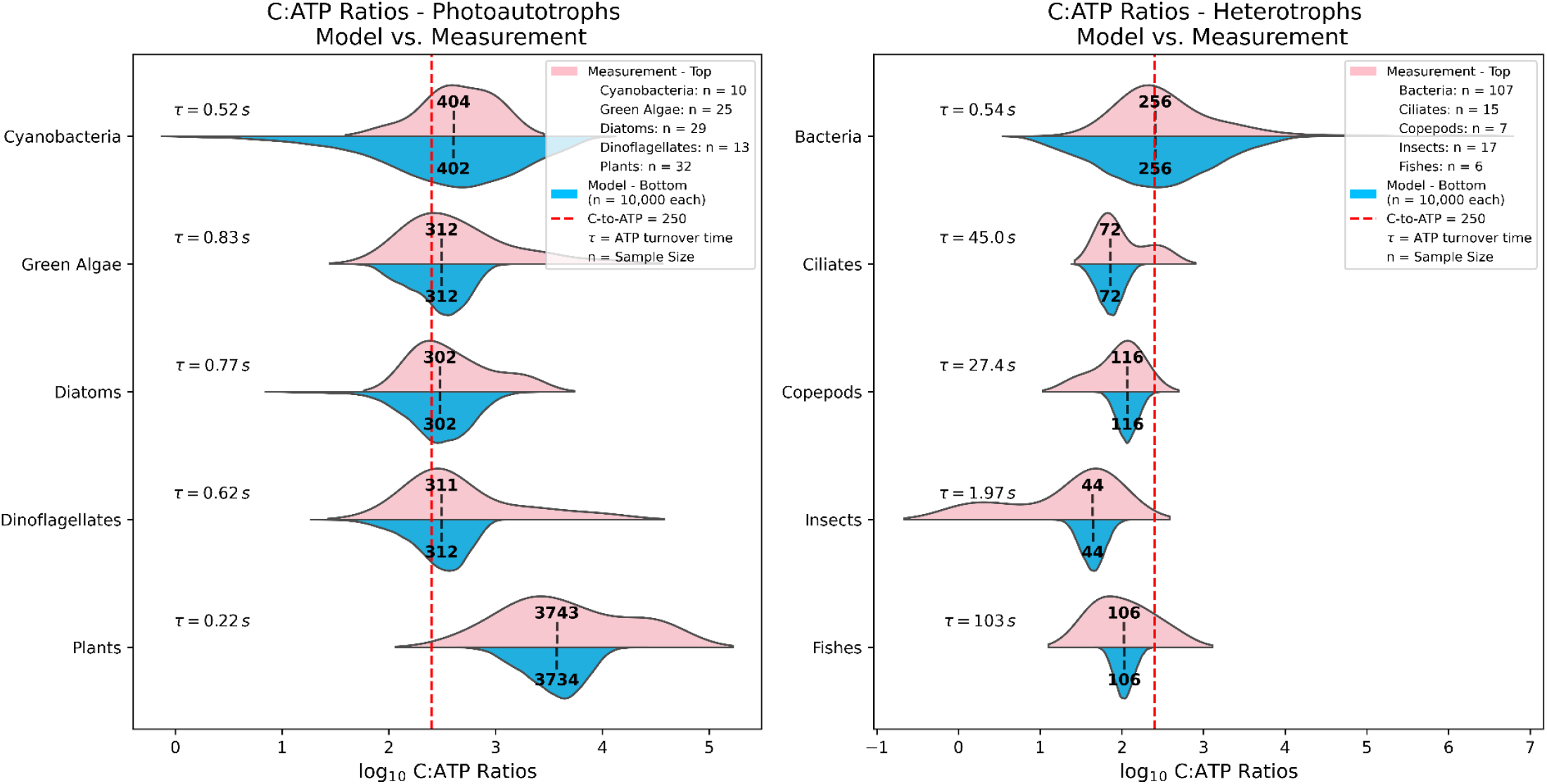
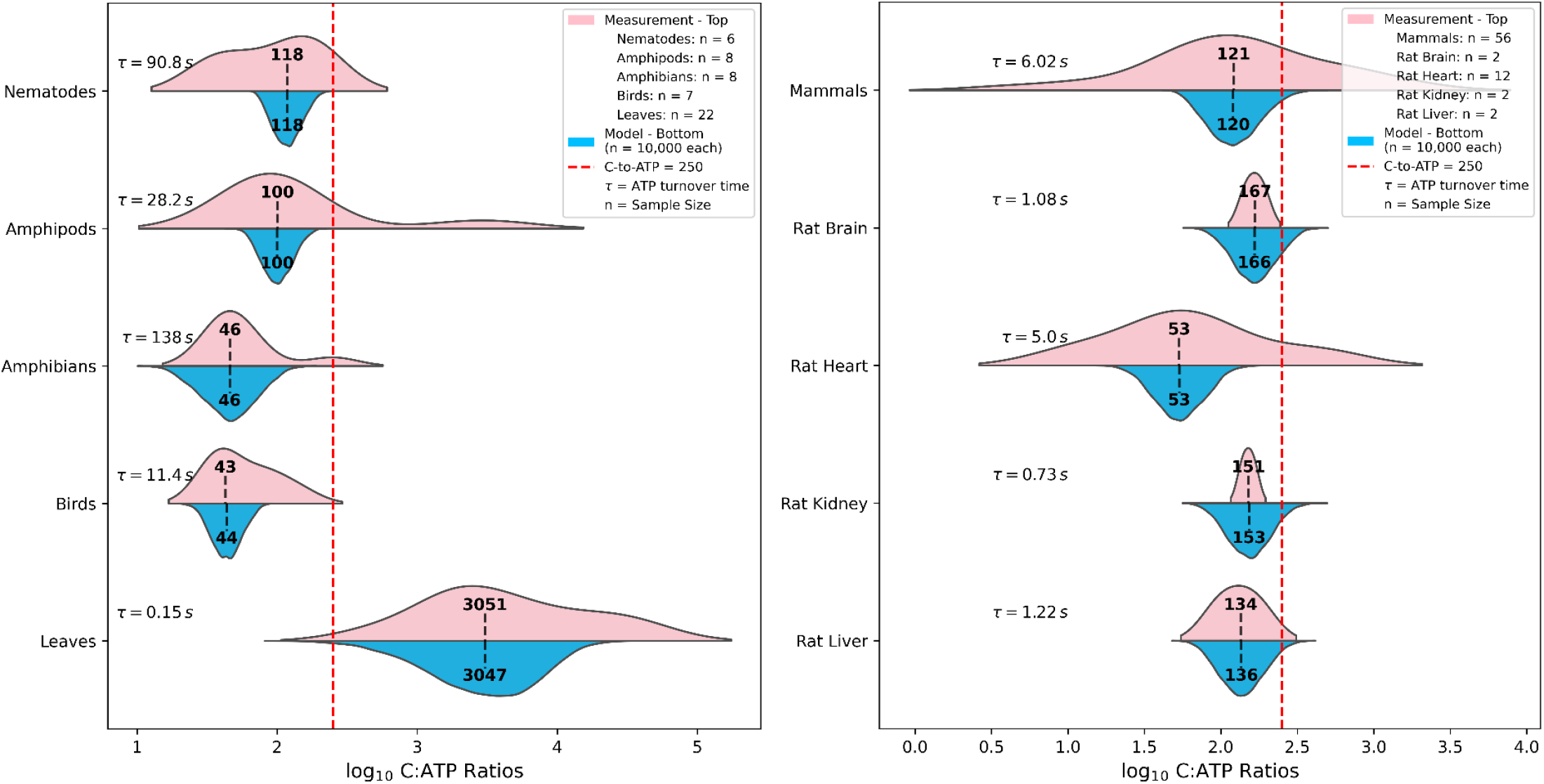
Comparison between experimental measurements and our theoretical steady-state model of the C:ATP ratio across major taxonomic groups, including unicellular and multicellular heterotrophs as well as photoautotrophs. For cyanobacteria, both data and model include N_2_-fixing and non-N_2_-fixing strains. Plant experimental ATP values are primarily from leaves, with some data from roots, shoots, and plant cell lines, whereas model values represent greenhouse-grown tree seedlings. For bacteria, both experimental and modeled values reflect fast-growing strains except for SAR11 in the data. Experimental data for ciliates represent tintinnids and Paramecium, whereas the model encompasses a broader range of ciliate types. Insects are represented by relatively large terrestrial species in both the data and the model; experimental ATP values are mostly from flight muscle, with some from brain and fat body. For fishes, ATP measurements are from white muscle and sperm cells. ATP data for amphibians are taken from sartorius muscle, gastrocnemius, and brain of frogs. ATP data for birds are taken from pectoral muscle of pigeons, starlings, domestic fowl, and pheasants, and from erythrocytes of chicken, pigeon, and turkey. ATP data for leaves represent various species, while the model represents lettuce. Mammalian ATP data are from a mix of tissues - including heart, brain, muscle, liver, kidney, thigh, fat pad, spleen, and blood - of rats, mice, pigs, dogs, and humans, as well as erythrocytes from twelve different animals. The rat tissue model uses respiration-rate data per tissue taken from [49], where brain, heart, kidney, and liver account for, respectively, 3%, 3%, 7%, and 20% of whole-body respiration. Details of the data compilation are provided in Supplementary Sections I and II, where all relevant references are listed. Densities are drawn using kernel density estimation from compiled literature data and model predictions with sensitivity analysis.

Heterotrophic bacteria are typically ∼1 μm in diameter, whereas photoautotrophic cyanobacteria reach ∼4 μm. Given these small cellular dimensions, the high copy number of ATP synthases in fast-growing aerobic bacteria, and the fact that ATP synthase complexes diffuse dynamically across the cytoplasmic membrane [42], we do not expect any substantial diffusion barrier for ATP molecule transport from its sites of production to sites of consumption. ATP diffusion times calculated from both the free-diffusion and target-search formulations (see Supplementary Section VI) fall well below one second. This is consistent with experimental data showing ATP turnover times of less than one second during exponential growth and nearly two seconds during basal growth in fast-growing bacteria [9,30].

Heterotrophic protists are, on average, substantially larger than bacteria and cyanobacteria, and most of their ATP is produced by mitochondria. Consequently, several factors become important determinants of ATP diffusion and thus ATP turnover time, including overall cell size, the number of mitochondria, their spatial distribution, the ATP consumption rate of different organelle types, and the intracellular packing density. A study of 13 heterotrophic protist species revealed that in cells larger than 10 μm, approximately 85% of mitochondria are located near the cell’s outer membrane [37], concluding that rapid access to oxygen is a key constraint shaping mitochondrial spatial distribution. Constructing an explicit model of ATP turnover time that incorporates all of these variables is beyond the scope of this paper, and free-diffusion and target-search formulations can provide only conceptual lower- and upper-bound estimates (see Supplementary Section VI). Thus, to obtain a more meaningful estimate of ATP turnover time in heterotrophic protists, we divide the empirical measurements of ATP content by our theoretical ATP production rates, resulting in an average turnover time of ∼45 seconds in ciliates. For comparison, the literature reports an upper bound of 30 seconds for ATP turnover time in *Paramecium tetraurelia* (a ciliate) [31].

Photoautotrophic protists are, on average, similar in size to heterotrophic protists, and their respiration rates are also comparable (see, for example, the respiration rates of ciliates and dinoflagellates in Table S5). However, two key differences distinguish these groups. First, chloroplasts can occupy up to 45% of the total cell volume in photoautotrophs [38,39] and produce their own ATP to satisfy local energy demands, reducing the volume of the cell that relies on mitochondrially produced ATP. Consequently, the effective respiration-to-volume ratio increases, which in turn lowers the ATP turnover time. The second major difference is that chloroplasts continuously generate oxygen via photosynthesis, creating an oxygen-rich intracellular environment. This enables mitochondria to position themselves close to chloroplasts, ensuring both immediate access to oxygen and proximity to other organelles, thereby minimizing ATP diffusion distances. Such mitochondrial positioning is consistent with three-dimensional reconstructions of intracellular organelle organization in microalgae [38]. Our comparison of the ATP content model with experimental ATP measurements supports these conclusions, indicating that ATP turnover times in microalgae are typically less than one second. ATP turnover time in higher plants is generally comparable to that observed in microalgae. As supported by our comparison between the ATP model and empirical ATP measurements, ATP turnover times in higher plants can still remain below one second. This is consistent with experiments reporting ATP turnover times below one second in *Arabidopsis* [30,32].

ATP turnover time in animals should be comparable to that in heterotrophic protists. Therefore, if one has data on the range and distribution of cell sizes that compose the body of an animal, it is possible to estimate the average ATP diffusion time. For instance, the size distribution and cell counts across all human tissues have recently been reported [36]. In adult males, individual cell mass ranges from 9.00 × 10^−12^ g to 7.23 × 10^−4^ g, with a weighted mean (excluding non-nucleated cells) of 5.83 × 10^−9^ g [36], corresponding roughly to a size range of 2.6 μm to 1114 μm, with a mean cell size of 22.3 μm. This average size is notably smaller than that of ciliates, typically reported around 100 μm [43]. This suggests that the average ATP turnover time in humans could potentially be shorter than in ciliates. However, another key factor must also be considered. Whole-body maximum respiration rates in animals vary widely from relatively low rates in fishes to significantly higher rates in terrestrial insects (see Figure S3 and Table S5). Although we earlier assumed that, under steady-state conditions, ATP production and consumption rates are equal, implying that respiration rate does not directly affect ATP turnover time, there is an important indirect effect. Consider two animals of the same size but with different respiration rates: the one with the higher rate will likely have a greater total mitochondrial volume or a higher mitochondrial population. Insects, for example, can have mitochondrial volumes reaching up to 43% of total cell volume [41,44], enabling mitochondria to be strategically positioned, with some near capillaries for optimal oxygen access and others near organelles for rapid ATP delivery. This mitochondrial arrangement partly explains why, according to our ATP model and its comparison with empirical ATP data, insects exhibit an ATP turnover time of approximately 2 seconds, while fishes, which have lower respiration rates, show turnover times around 103 seconds. Additionally, in animals, oxygen diffusion from the circulatory system (e.g., blood vessels) to individual cells presents another constraint on ATP production rate. For further discussion, see Supplementary Section VI.

### 2.2. Carbon Content

Carbon content 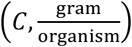 ranges from as low as 5% of dry mass in ctenophores [45] to as high as 65% in the leaves and stems of plants [46], but generally falls between 40% and 50% of an organism’s dry mass. Given that carbon turnover time is much longer than ATP turnover time, meaning carbon content remains relatively stable over timescales during which ATP levels fluctuate across different physiological states, we estimate carbon content using established equations that relate organism’s dry mass 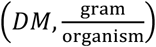 to carbon content (see Supplementary Section II). Structural materials 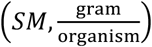, such as bacterial cell walls or vertebrate skeletons, can be either metabolically active (during synthesis, remodeling, degradation, motility, etc.) or inactive depending on context. Since our goal is to estimate metabolically active organismal carbon content, we partially exclude *SM* from the organism’s *DM* before calculating carbon content. To do this, we use experimentally measured values of the maximum *SM* mass relative to *DM* obtained from the literature (see Table S4 and Figure S5). We then consider a range of *SM* values, from zero, where the entire organismal mass is metabolically active, to the measured maximum, where *SM* is entirely inactive and should be excluded from *DM*. The equation used to calculate carbon content, *C*, is:

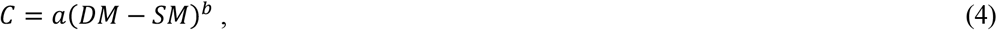

where *a* and *b* represent the scaling coefficient and scaling exponent, respectively.

## 3. Results and Discussion

Experimental measurements closely align with our model predictions across a wide range of photoautotrophs (cyanobacteria, green algae, diatoms, dinoflagellates, and plants), heterotrophs (bacteria, ciliates, copepods, amphipods, nematodes, insects, fishes, amphibians, birds, and mammals), and tissues, with good agreement in both the median C:ATP values for each group as well as the variation range within groups (Fig. 1).

Moving through the taxonomic ranks, prokaryotes have a median C:ATP of 256, but exhibit extremely wide variation - from as low as 28.4 in exponentially growing *Streptococcus bovis* to as high as 1,800,000 in SAR11 after 20 h of starvation followed by 2 h of 2-propanol treatment (SAR11 is a slow-growing bacterium and one of the most abundant species in surface ocean waters [47]) (Fig. 1 and Supplementary Section I). Eukaryotes show a broad range of median C:ATP values across groups and tissues, from as low as 43 in birds to as high as 3,743 in plants. Within each group, the variation is even more substantial: C:ATP ranges from 1.2 in the brain of *Helicoverpa armigera* during the non-diapausing pupal stage to 47,801 in a deep-sea sponge. The major differences between prokaryotes and eukaryotes arise from: (1) the exceptionally strong size dependence of respiration rates in fast-growing prokaryotes with a scaling exponent near 2 for heterotrophic bacteria, making respiration far more sensitive to cell-size variation than in eukaryotes (this size dependence is further reflected in the normalized ATP content data, see Supplementary Section I); and (2) the presence of mitochondria and chloroplasts in eukaryotes, combined with their broader range of cell sizes, which together shape the diverse ATP turnover times observed across eukaryotic groups.

The C:ATP ratio is, on average, higher in eukaryotic photoautotrophs than in eukaryotic heterotrophs, largely due to the close *spatial* coupling of chloroplasts, mitochondria, and other organelles, which facilitates rapid oxygen access and short ATP turnover times. In prokaryotes, the C:ATP ratio in photoautotrophic cyanobacteria is higher than in heterotrophic bacteria primarily because fast-growing heterotrophic bacteria have substantially higher respiration rates than cyanobacteria at comparable cell sizes. Our dataset also indicates that nitrogen-fixing cyanobacteria have, on average, a lower C:ATP ratio than non-nitrogen-fixing strains, consistent with their elevated respiration rates required to meet the ATP demands of nitrogen fixation. Among photoautotrophs, plants exhibit substantially higher C:ATP ratios than microalgae (Figure 1). This arises from their much larger cell-size range [48], which yields diverse ATP turnover times across cells, different respiratory demands, and a high proportion of structural biomass - cell walls alone can constitute more than 90% of plant dry mass. As shown in Figure 1, our model predicts that plants have, on average, an ATP turnover time of about threefold lower than that of microalgae.

Diatoms and dinoflagellates have generally similar C:ATP ratios, although diatoms tend to have slightly lower values. Several factors contribute to this difference. First, diatoms have a slightly larger median size than dinoflagellates. They also exhibit lower respiration rates (by a factor of 3-4 for median-sized species), higher photosynthesis rates (approximately twofold on average), and greater structural material content (also about twofold higher on average). Together, these factors suggest that diatoms have a slightly lower C:ATP ratio than dinoflagellates.

Most C:ATP values reported for animal groups do not represent whole-body C:ATP; instead, they reflect tissue-specific C:ATP, which varies with the metabolic activity of each tissue - and that activity differs widely among species. For example, in rats the ratio of “% body oxygen use” to “% body mass” ranges from 0.71 for skeletal muscle (30% O_2_ use / 42% mass) to 7.78 for the kidney (7% O_2_ use / 0.9% mass), spanning more than an order of magnitude [49]. In humans the pattern shifts: skeletal muscle has the lowest ratio (0.48), whereas the heart has the highest (27.5), a >50-fold difference [49] (See Supplementary Section II.4.2 for model implementation details). Beyond organ-level respiration rates, ATP turnover time also varies substantially across tissues, as shown in Fig. 1 and the Supplementary Section VI. ATP turnover time depends on multiple factors, including the type of circulatory system, the distance between blood vessels and target cells, the distribution of cell sizes within the tissue, and the mitochondrial population and spatial organization.

Holm-Hansen, the pioneer of the “universal C:ATP” concept, reported from 30 algal cultures that the ratio was independent of cell size [50]. In contrast, using our compiled dataset spanning a broad range of organisms (see Supplementary Section I), we observed statistically significant size-related patterns in C:ATP for bacteria, microalgae, ciliates, insects, amphibians, birds, and mammals. Importantly, these correlations differ in sign: in some groups C:ATP decreases with size, while in others it increases. Our current theoretical framework is a size-dependent model in all respects except for ATP turnover time, which we treat as a species-specific constant in our analysis. Understanding the origin of size-dependent C:ATP patterns requires a size-dependent model of ATP turnover time, which in turn would need data on cell sizes, mitochondrial abundance and spatial organization, intracellular packing density, and ATP consumption rates of different organelles. Developing such a model is beyond the scope of this paper. Given this limitation, we propose that the observed patterns of C:ATP versus organism or tissue size reflect the balance between two scaling relationships: (i) respiration rate versus size and (ii) carbon content versus size (see Supplementary Section VII). For example, Table S5 shows that in bacteria the scaling exponent for respiration rate versus dry mass is 1.96, whereas the exponent for carbon content versus dry mass is 1.22. Because respiration increases more steeply with size than carbon content, C:ATP should decrease with increasing bacterial cell size - precisely matching the trend observed in our compiled dataset.

Experimental data show substantial variation in C:ATP across multiple groups, reflected in the skewed distributions (Fig. 1). Here we highlight a few representative examples; full details of the physiological and environmental conditions under which ATP was measured are provided in Supplementary Section I. In insects, the left-tailed distribution is driven by the high ATP content of the cotton bollworm brain during the pupal stage, illustrating how ATP levels change markedly across developmental stages. In amphipods, the right-tailed distribution is caused by the low ATP content of *Gammarus lacustris* exposed to gradual warming in saltwater at 9 °C. Among mammals, the highest ATP content is reported in epididymal adipose tissue of rats, whereas the lowest is found in horse erythrocytes.

The C:ATP ratio may vary further under non-equilibrium conditions, which are not considered in the present work. Studies show that ATP content fluctuates with growth phase, carbon source, and cell cycle dynamics [51,52]. For instance, *E. coli* and *P. putida* exhibit transient ATP accumulation during the shift from exponential to basal growth, due to a faster drop in ATP consumption than production especially pronounced with acetate or oleate as carbon sources [51]. Single-cell studies reveal that ATP levels in *E. coli* can fluctuate 1.5-to 2-fold during exponential growth, with more stable ATP in fast-growing cells and greater fluctuations in slower-growing ones [52]. These changes are tied to the cell cycle, mitochondrial dynamics, and cyclin-dependent kinase activity [53,54].

Patel *et al*. [55] proposed that ATP acts as a hydrotrope (in addition to its role as the universal energy currency), requiring millimolar concentrations to keep proteins soluble despite enzymes needing only micromolar levels. Bochdansky *et al*. [6] noted that this explains the universally conserved cytoplasmic ATP concentration, making it a reliable proxy for microbial cytoplasmic volume and biomass in ocean studies. Aside from the proposition that ATP may act as a hydrotrope (itself still debated [56]), the arguments above face several criticisms. First, as we document in this paper, the normalized ATP concentration is not universally millimolar but instead varies widely across the tree of life (see the normalized ATP content compiled in Supplementary Section I showing variation by 6-orders of magnitude). Second, the micromolar ATP requirements of enzymes cannot account for the total rate of ATP consumption, because a major fraction of cellular energy is dissipated as heat due to the inefficiency of intracellular machineries [57]. The actual rate of ATP consumption at the level of cellular traits and organelles remains an active field of study with many unknowns [2]. Third, as we argue here, the accumulation of ATP in cells is best explained by ATP turnover time, which varies extensively across organisms and depends at least on cell size, mitochondrial abundance and spatial distribution, intracellular packing density, and the ATP consumption rate per organelle types.

## Acknowledgement

This work was supported by the Simons Collaboration on Computational Biogeochemical Modeling of Marine Ecosystems/CBIOMES (Grant Id: 549937, Z.V.F. and 549935, A.J.I.). The authors express their gratitude to Dr. Manon Laget, Dr. Dany Crouteau, Dr. Niall McGinty, and Dr. Ruby Hu (all from Dalhousie University) for their helpful discussions.

## Supplementary Information

## Supplementary Methods

### I. Compiled measurements of ATP content per unit biomass across the kingdoms of life

We compiled 397 ATP-content measurements from 82 papers published between 1964 and 2024, spanning a wide diversity of taxa. The dataset encompasses bacteria (9 classes, n = 108 data points), microalgae (13 classes, n = 113), heterotrophic protists (1 class, n = 16), arthropods (3 classes, n = 24), Mollusca and nematodes (3 classes, n = 9), insects (n = 17), amphibians (n = 8), fish (2 classes, n = 6), birds (n = 7), mammals (n = 56), sponges (n = 1), and plants (n = 32). Across animal and plant tissues, the data include muscle (n = 34), brain (n = 11), heart (n = 17), kidney (n = 2), liver (n = 5), fat body (n = 5), blood and red blood cells (n = 20), spleen (n = 1), sperm (n = 3), leaves (n = 22), and roots, shoots, and other plant cell lines (n = 10). All data are provided in the Supplemental Excel File 1.

For each study, the file includes: (1) reference; (2) ATP assay method; (3) physiological factors affecting ATP content; (4) class; (5) group; (6) common name/species; (7) tissue group; (8) physiological state for which ATP content is reported; (9) original ATP units; (10) ATP content in original units; (11) ATP in grams per gram of dry mass; (12) tissue or cell dry mass (g); (13) carbon content (g); (14) ATP per cell or tissue (g); (15) C:ATP ratio; and (16) method used to estimate tissue or cell dry mass.

For every species or tissue, we report the minimum and maximum ATP content in the original units, followed by conversion to a common unit (grams ATP per gram dry mass) to allow comparison across studies. As illustrated in Figure S1, normalized ATP content spans 2.78×10^−7^ to 4.46×10^−1^ g ATP g^−1^ dry mass, representing over six orders of magnitude variation.

Because most studies did not report total organism or organ mass, we estimated masses using several approaches described in Supplementary Excel File 1 (see the column “How the tissue or cell dry weight is estimated?”). For example, in mammals, organ masses were estimated using published scaling relationships that relate whole-organism mass to organ-specific mass. These estimates allowed us to evaluate how the C:ATP ratio scales with organism or tissue dry mass. As shown in Figure S2 (detailed version given in Supplementary Excel File 2), the C:ATP ratio exhibits statistically significant size-dependent patterns in several groups, based on the p-values of the regression slopes of log_10_(C:ATP) versus log_10_(dry mass). In bacteria, microalgae, birds, and mammals, the correlation is negative, whereas in insects, ciliates, and amphibians, it is positive. Across tissues, statistically significant patterns are observed in erythrocytes and leaves (negative correlation) and in brain tissue (positive correlation).

**Figure S1:**
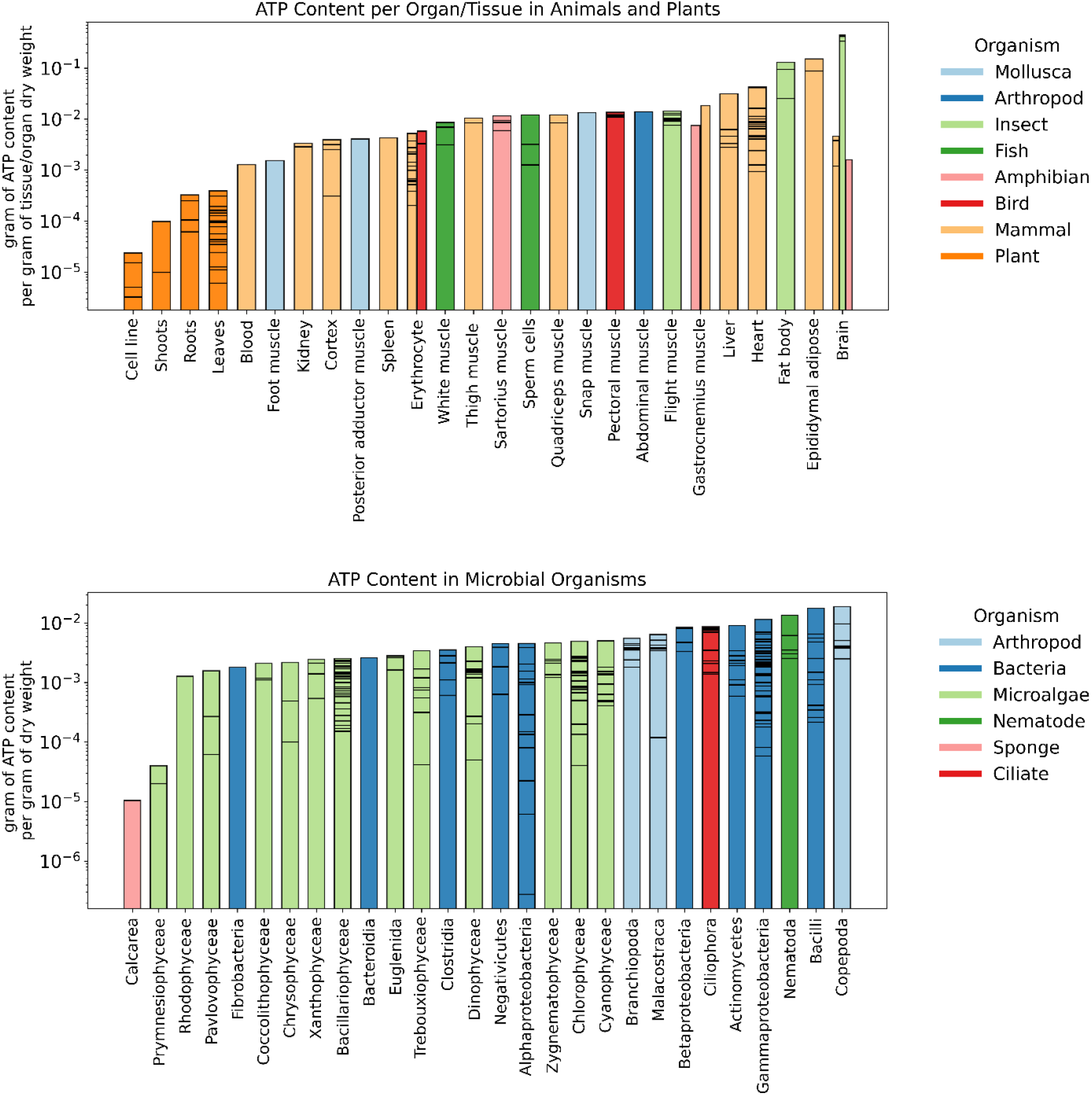
ATP content (g per g of dry mass) in various organs across different animal groups (top panel, n = 130 data points) and across diverse microbial organisms (bottom panel, n = 265 data points) under different physiological and environmental conditions. Each color represents a specific animal or microbial group, as indicated in the legends. Horizontal lines within the bars indicate ATP values of individual organisms within the same group.

**Figure S2:**
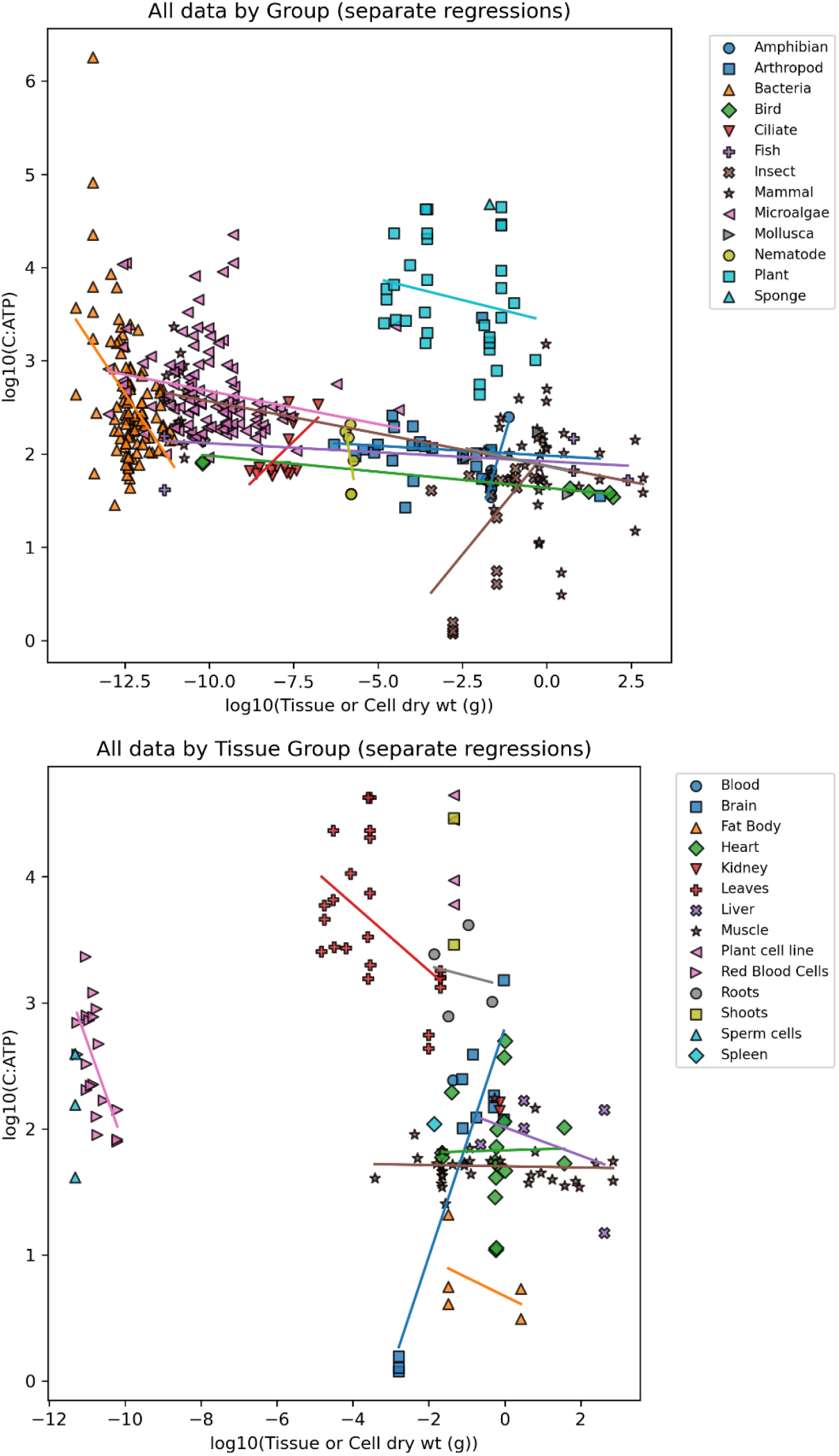
Size-dependence of the C:ATP ratio across organisms and tissues. Statistically significant slopes show decreasing C:ATP with mass in bacteria, microalgae, birds, mammals, erythrocytes, and leaves, and increasing C:ATP with mass in insects, ciliates, amphibians, and brain tissue.

### II. Data Compilation from the Literature for the Theoretical Model

To estimate C:ATP ratios, we assembled dry-mass ranges (Table S1) and collated equations predicting respiration rate, carbon content, and structural material composition based on organism size (Tables S2, S3, and S4), then we standardized the units for all variables (Table S5) across a diverse set of organisms including bacteria, microalgae, heterotrophic protists, multicellular zooplankton, freshwater invertebrates, terrestrial insects, nematodes, fishes, amphibians, birds, mammals, and vascular plants. The procedure used to perform the standardization is described in detail in Supplementary Section II.4. To study the distribution around the median C:ATP ratio, we performed a sensitivity analysis for all parameters with a range of values. We randomly selected 10,000 samples using a normal distribution centered around the mean or median value, bounded by the minimum and maximum observed values. If the mean or median was unavailable, we used a uniform distribution instead. We compared the results of our C:ATP model with experimental measurements (∼400 data points from 82 papers published between 1964 and 2024; Supplementary Excel File 1).

#### II.1. Range of Dry Mass in Studied organisms

**Table S1:**
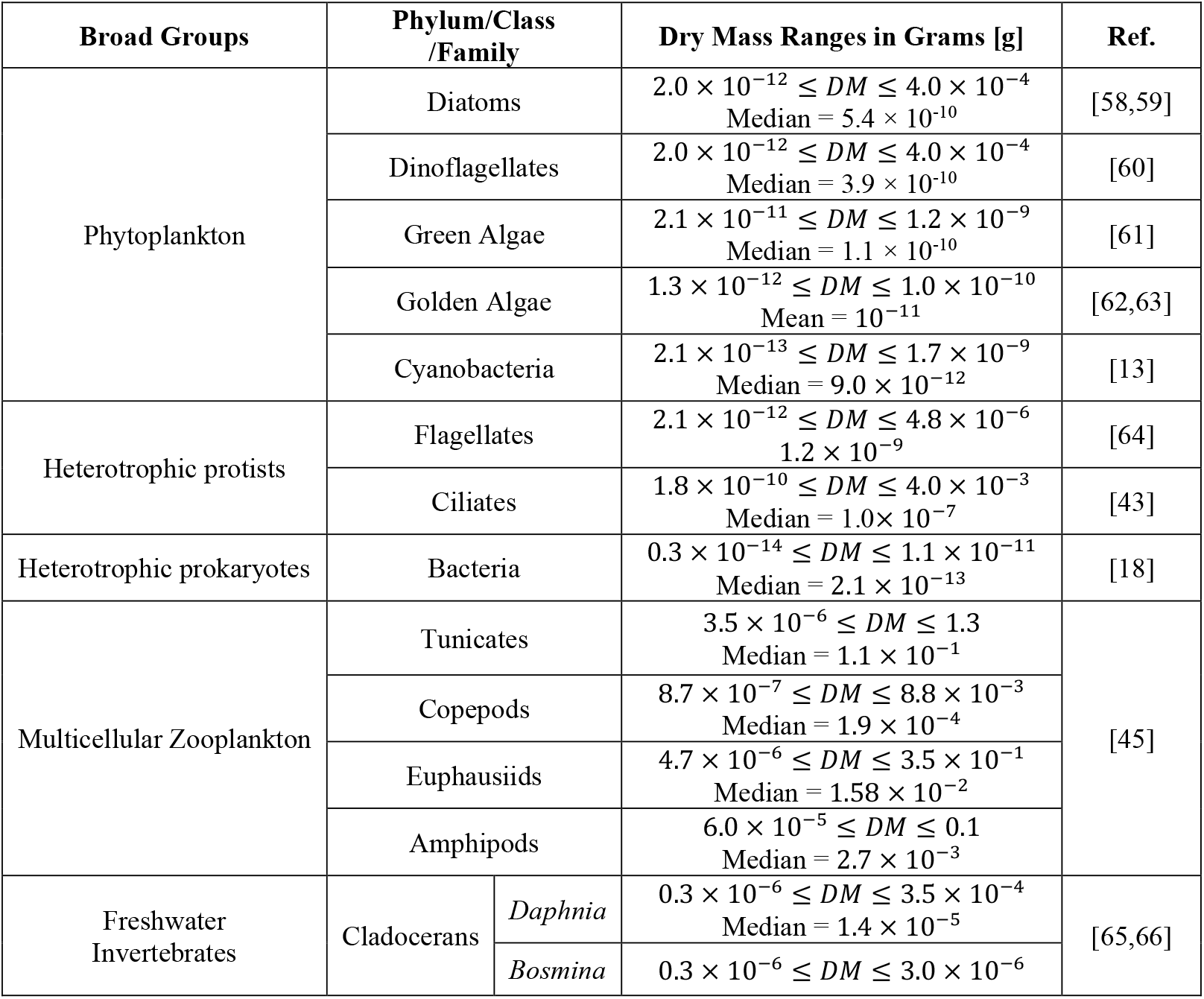
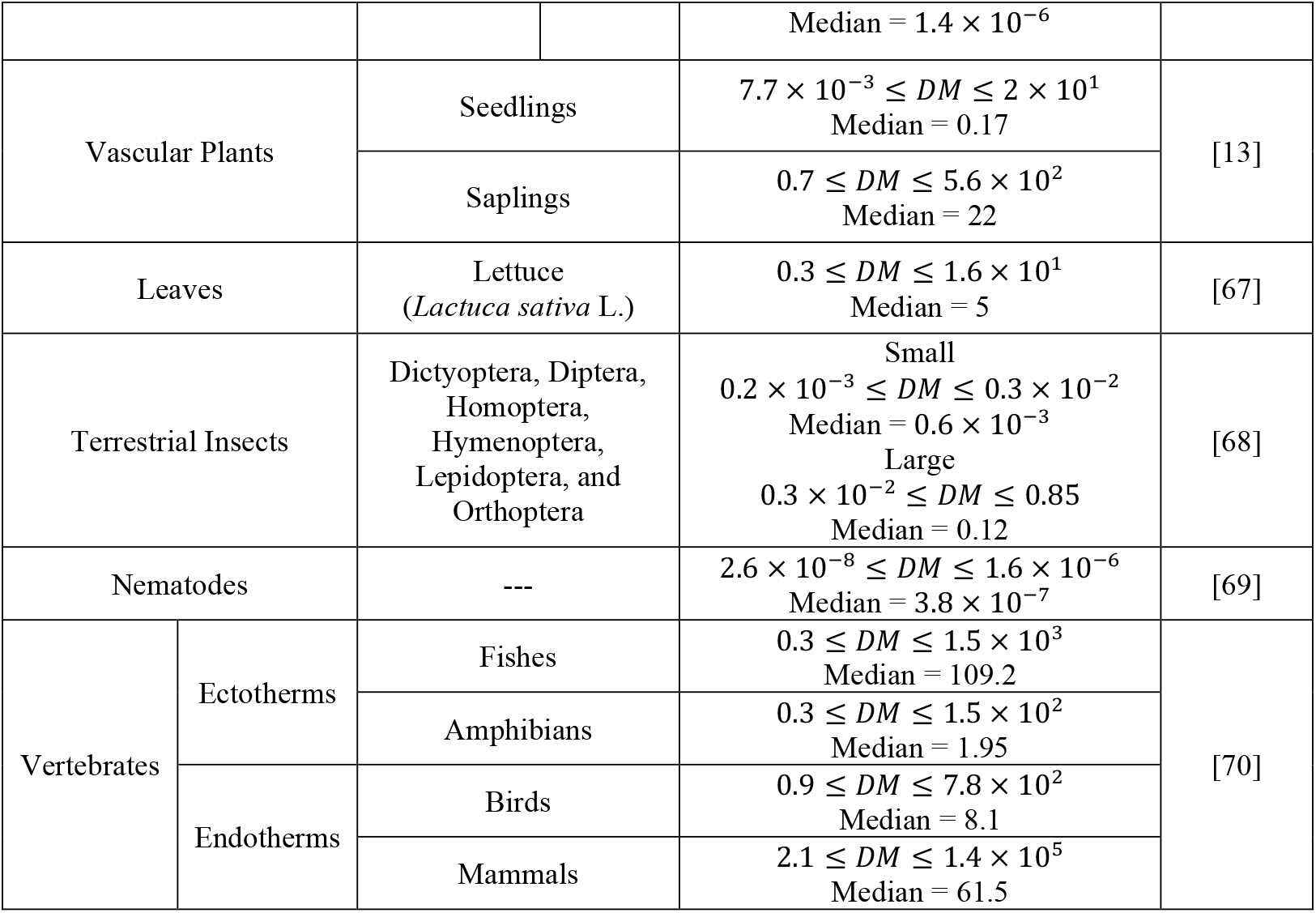
Range of dry mass for the studied organisms, extracted directly from the references or calculated assuming a dry mass-to-body mass ratio of 0.3 and a mass density similar to that of water.

#### II.2. Aerobic Respiration Rate

We have compiled empirical scaling equations from various sources that relate respiration rates in heterotrophs and dark respiration rates in photoautotrophs to organism size (Table S2). These equations apply to organisms in an actively growing state for protists and prokaryotes, food-saturated growth rates for multicellular zooplankton, maximum growth rates for freshwater invertebrates, maximum metabolic rates for vertebrates and terrestrial insects, a naturally growing state for field tree saplings, and growth in nitrogen-enriched soil for greenhouse tree seedlings. We retain the original units provided by each reference in Table S2 but standardize all units to a unified format in Table S5. Additionally, we convert all the original temperatures to 25°C using a Q_10_ value of 2.5 [71]. For endothermic vertebrates, we fix the temperature at 38°C. Where data or equations relating organism volume to dry mass are unavailable, we use the relationship *DM*[pg] ≅ 0.57(*V*[μm^3^])^0.92^, as developed by Lynch [1]. If equations are provided in terms of body mass, we assume that dry mass is 30% of the body mass [13]. Figure S2 depicts the respiration rate in both heterotrophs and photoautotrophs as a function of dry mass, illustrating how respiration rate varies across species. The dry mass in the figure spans 20 orders of magnitude.

**Figure S3:**
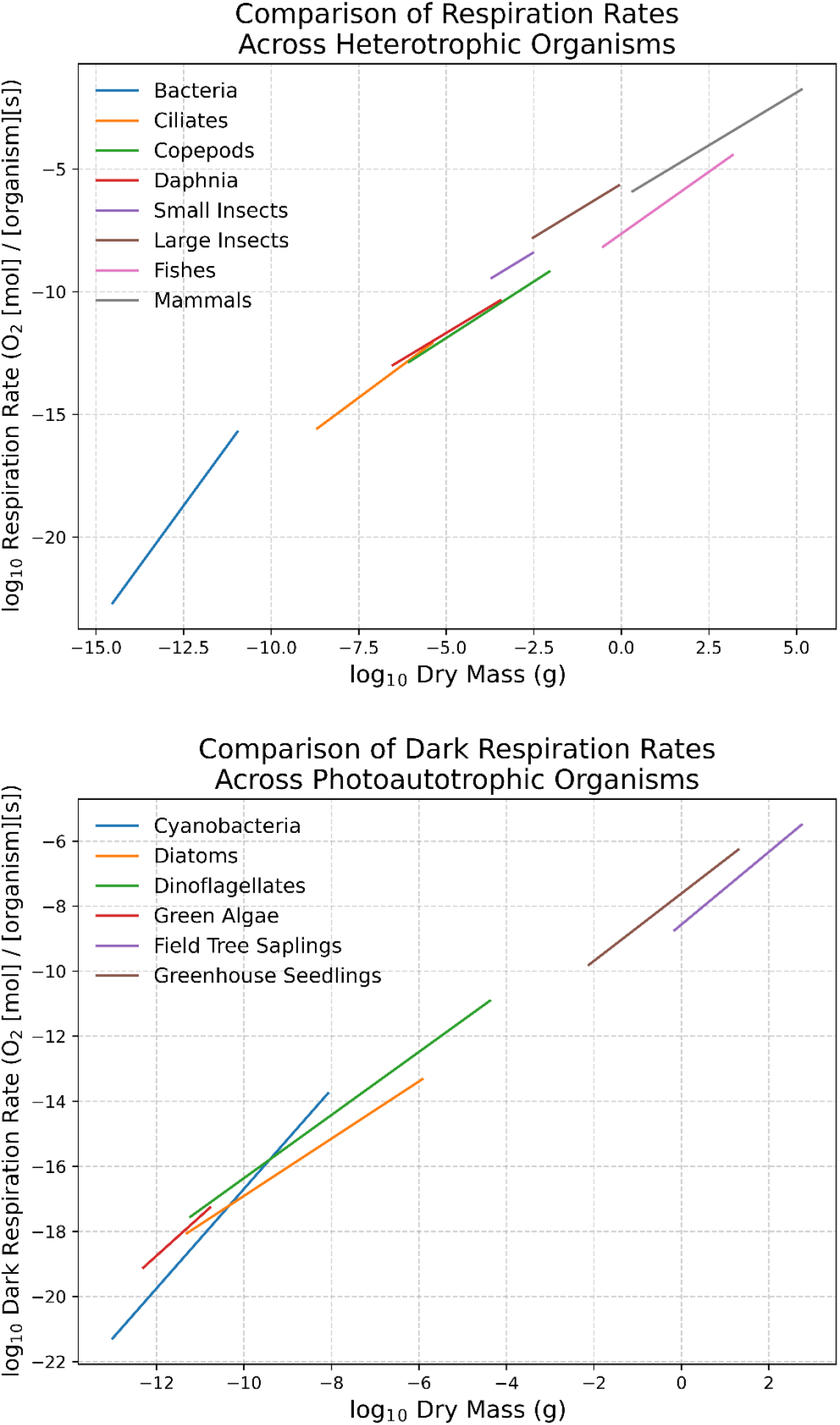
Respiration rate in heterotrophic (top panel) and photoautotrophic (bottom panel) organisms as a function of dry mass, ranging from bacteria to mammals and spanning 20 orders of magnitude. The respiration rates represent organisms in their active state.

**Table S2:**
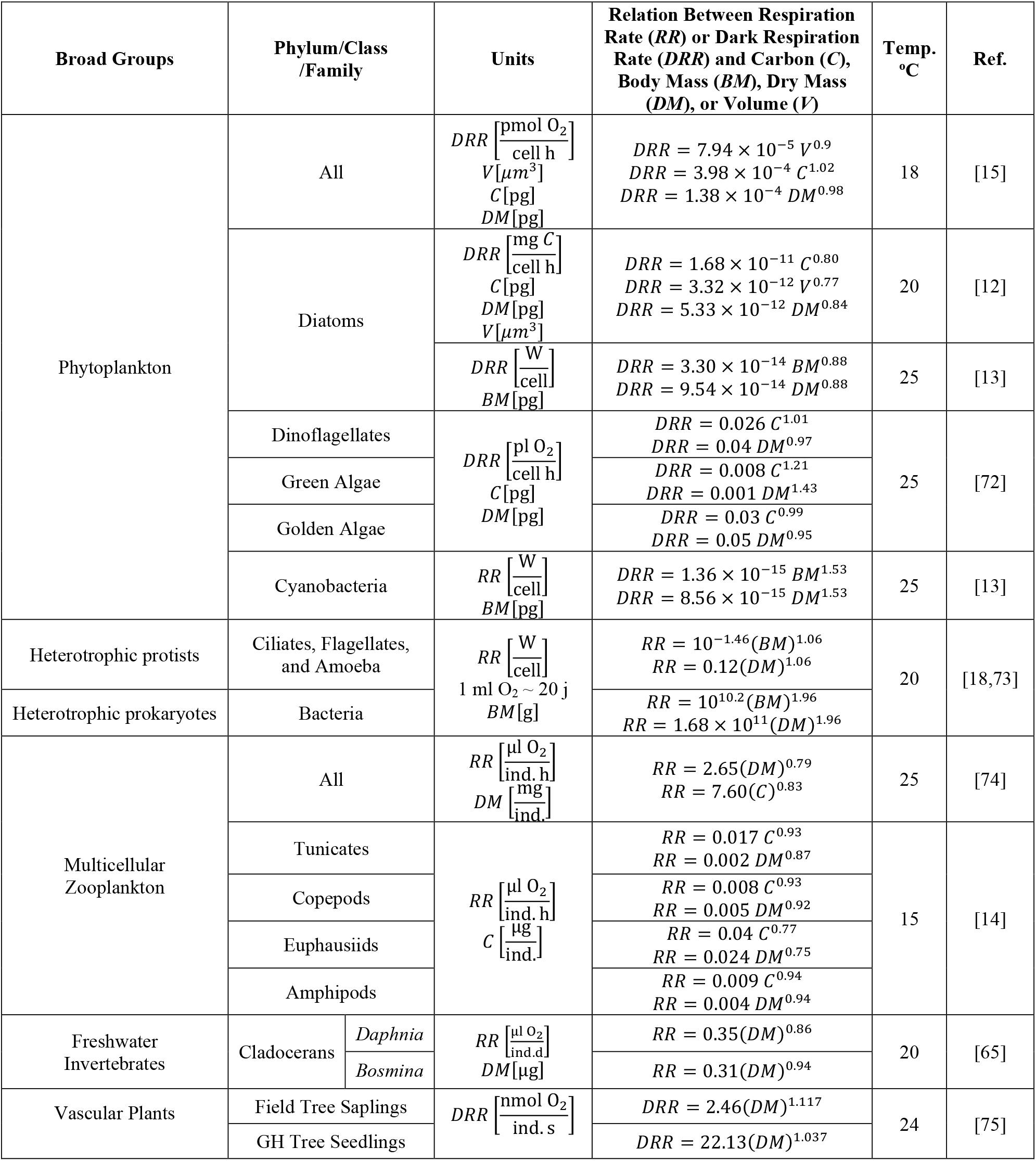
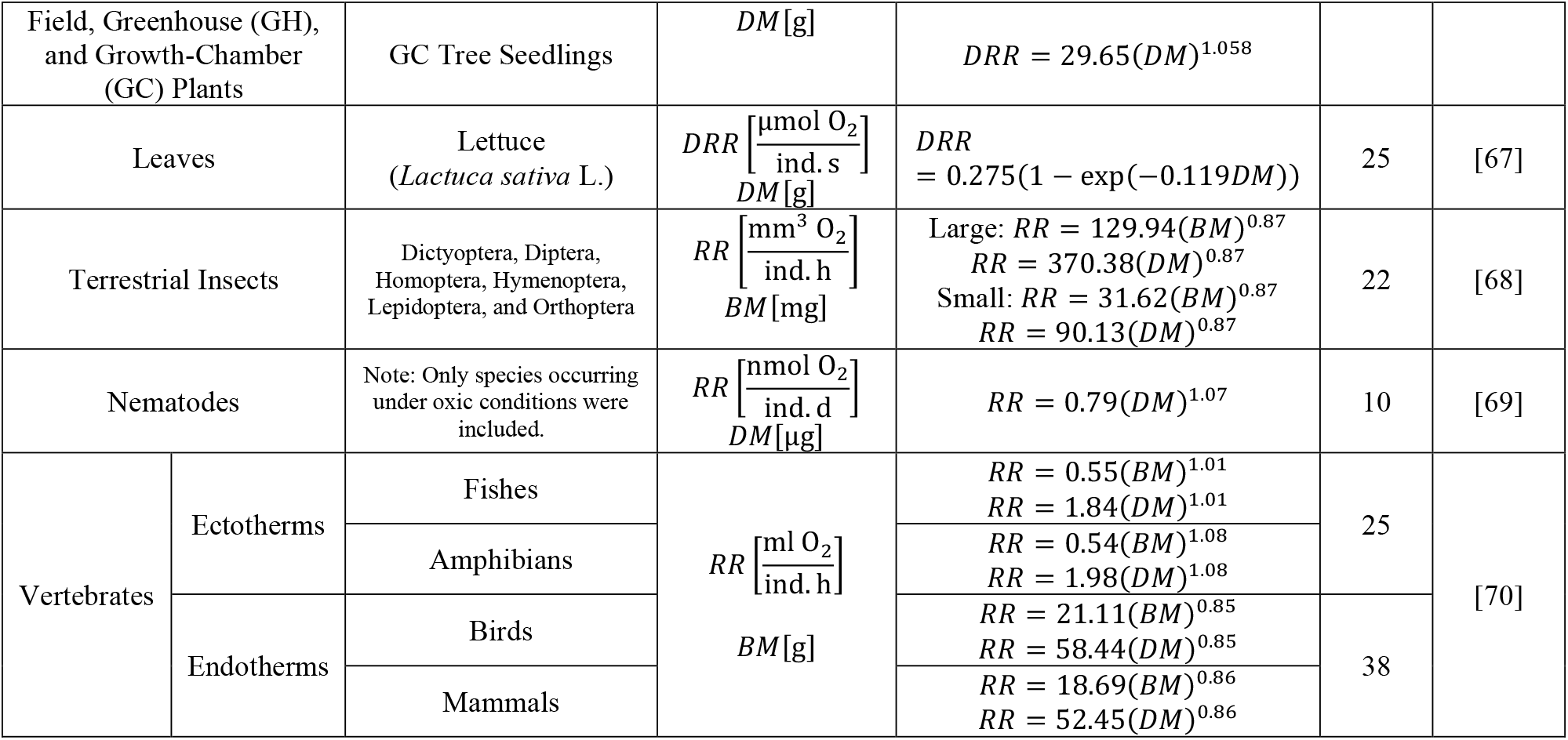
Mass scaling of respiration rates across the kingdoms of life. For details on how these equations were derived, see the section “Standardization of Respiration Rates and Carbon Content Across Units and Temperature” in the Supplementary Section II.4.

#### II.3. Carbon Content Across the Kingdoms of Life

Carbon turnover time is significantly higher than ATP turnover time, meaning carbon content remains stable over timescales in which ATP content fluctuates. This stability allows us to compute carbon content from dry mass using power-law functions. This approach is reliable due to the strong correlation observed, as evidenced by the high R^2^ values in the empirical equations (Table S3). An alternative method estimates carbon content using maximum growth rate scaling equations. While this method offers insights into carbon content increase per unit time, it typically produces lower R^2^ values, ranging from as low as 0.001 in larval fish to 0.733 in crustaceans, with other organisms falling between these extremes [66]. Consequently, we have compiled data on the mass-scaling of carbon content per organism from various sources, with the summarized results presented in Table S3. The same equations, standardized to consistent units, are provided in Table S5. Figure S4 compares the carbon content versus dry mass across different organisms, showing those with a known power-law relationship (top panel) and those for which only the carbon-to-dry-mass ratio is available (bottom panel).

**Figure S4:**
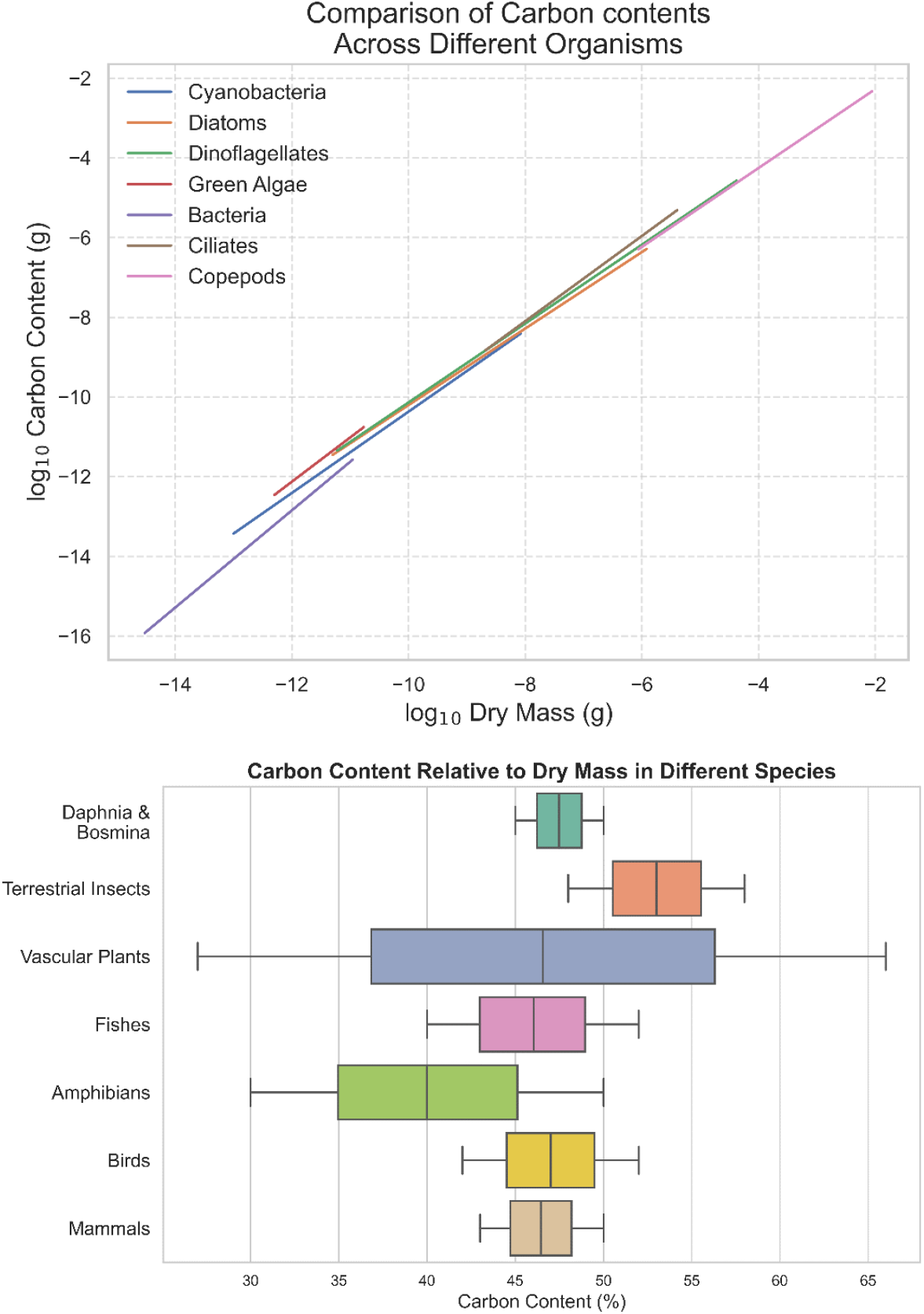
Carbon content versus dry mass relationship across different organisms (top and bottom panels).

The literature suggests that structural materials (SM) such as frustules, theca, lorica, coccoliths, siliceous scales, cell walls, tunics, chitin, shells, molts, and skeletons should be excluded from dry mass before carbon calculation because these components are metabolically inert once formed. While this is partially correct, these materials (some composed of inorganic substances and others of organic compounds) can actively contribute to metabolic processes during their synthesis, remodeling, degradation, and motility. As an example of the metabolic activity of structural materials, microorganisms under turbulent conditions undergo dynamic shape changes that alter their centers of buoyancy, mass, and hydrodynamic stress [76]. These changes enable them to adjust their orientation and escape turbulent currents, as observed in dinoflagellates and raphidophytes [76]. These adaptations can not be made possible without the dynamic properties of their structural materials.

To evaluate the impact of SMs on C:ATP ratios, we treat them as variably contributing to metabolically active dry mass. Rather than assuming a fixed exclusion, we implement a bounded uncertainty approach in which the structural fraction of dry mass can vary continuously between zero and an upper limit derived from the literature. Specifically, the maximum reported contribution of SMs to organism dry mass (summarized in Figure S5 and Table S4) is used to define the upper bound of this range, ensuring that all empirically observed cases are included.

In the simulations, the fraction of dry mass allocated to SM is randomly sampled from a uniform distribution between 0 and this maximum value, reflecting the fact that SM may be metabolically active under some conditions and largely inactive under others. The metabolically active dry mass entering the C:ATP calculation is then computed by subtracting this sampled structural fraction from total dry mass. This adjusted dry mass is used in Eq. (4) to estimate carbon content.

**Figure S5:**
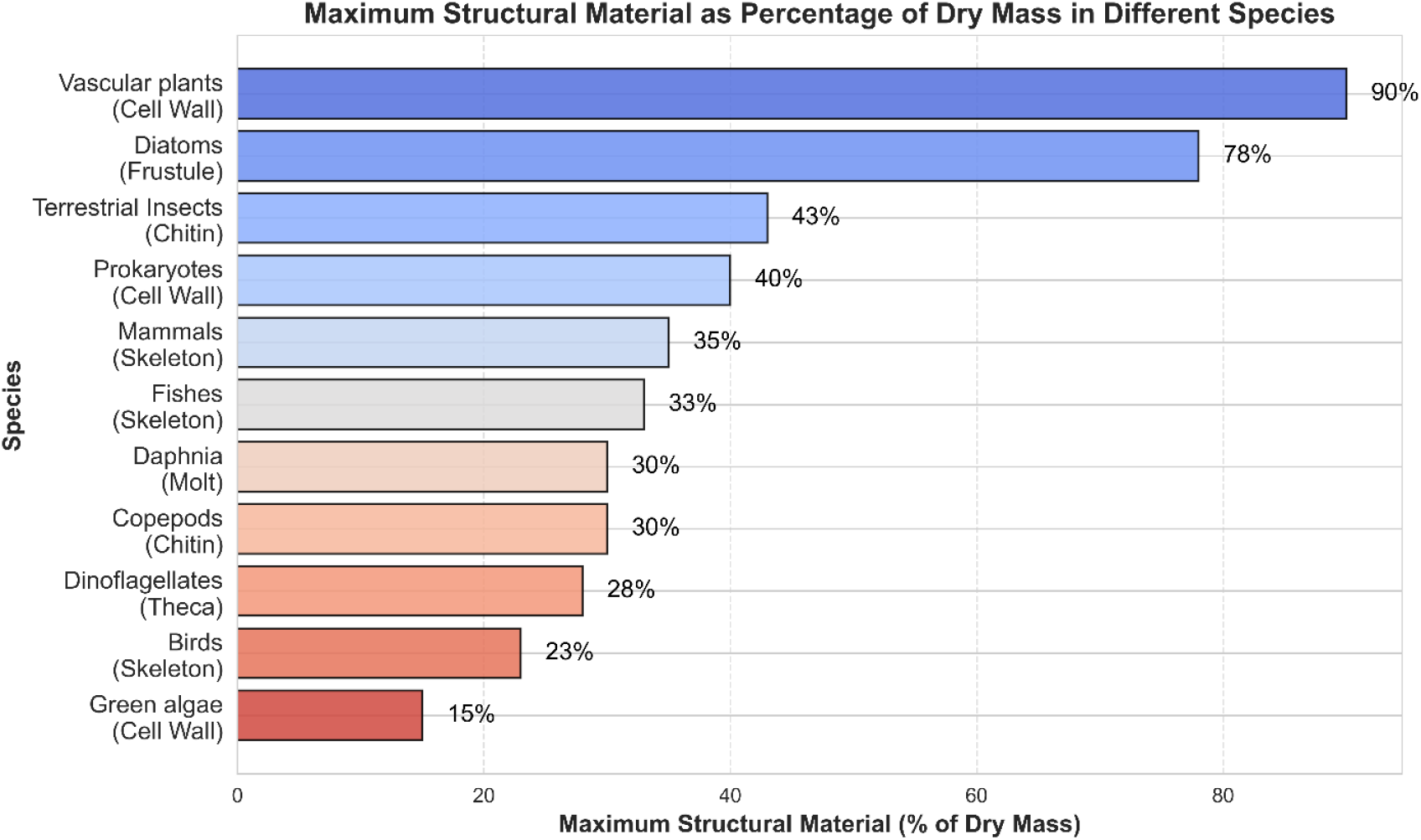
Percentage of maximum structural material relative to dry mass across different taxa.

The exclusion of storage oils or lipid droplets, referred to as non-living carbon, from carbon estimations has been proposed [77]. However, while lipid storage may sometimes be temporarily metabolically inactive, it remains highly dynamic and potentially active, contributing to numerous cellular processes. Recent research has highlighted their roles in metabolism, energy homeostasis, signaling, and buoyancy regulation, among others [78–81]. In eukaryotes, lipid droplets establish active interactions with organelles such as the endoplasmic reticulum, mitochondria, Golgi apparatus, and vacuoles, which are essential for the normal cellular life cycle [78]. The copy number and size of these droplets also respond dynamically to the type of food (carbon source) and changes in cell volume [40]. In prokaryotes, lipids like poly(3-hydroxybutyrate), polyhydroxyalkanoates, triacylglycerols, and wax esters play active roles in responding to stress, imbalanced growth, and starvation conditions [80]. Hence, we include lipid storage mass in our estimates of carbon-to-ATP ratios.

**Table S3:**
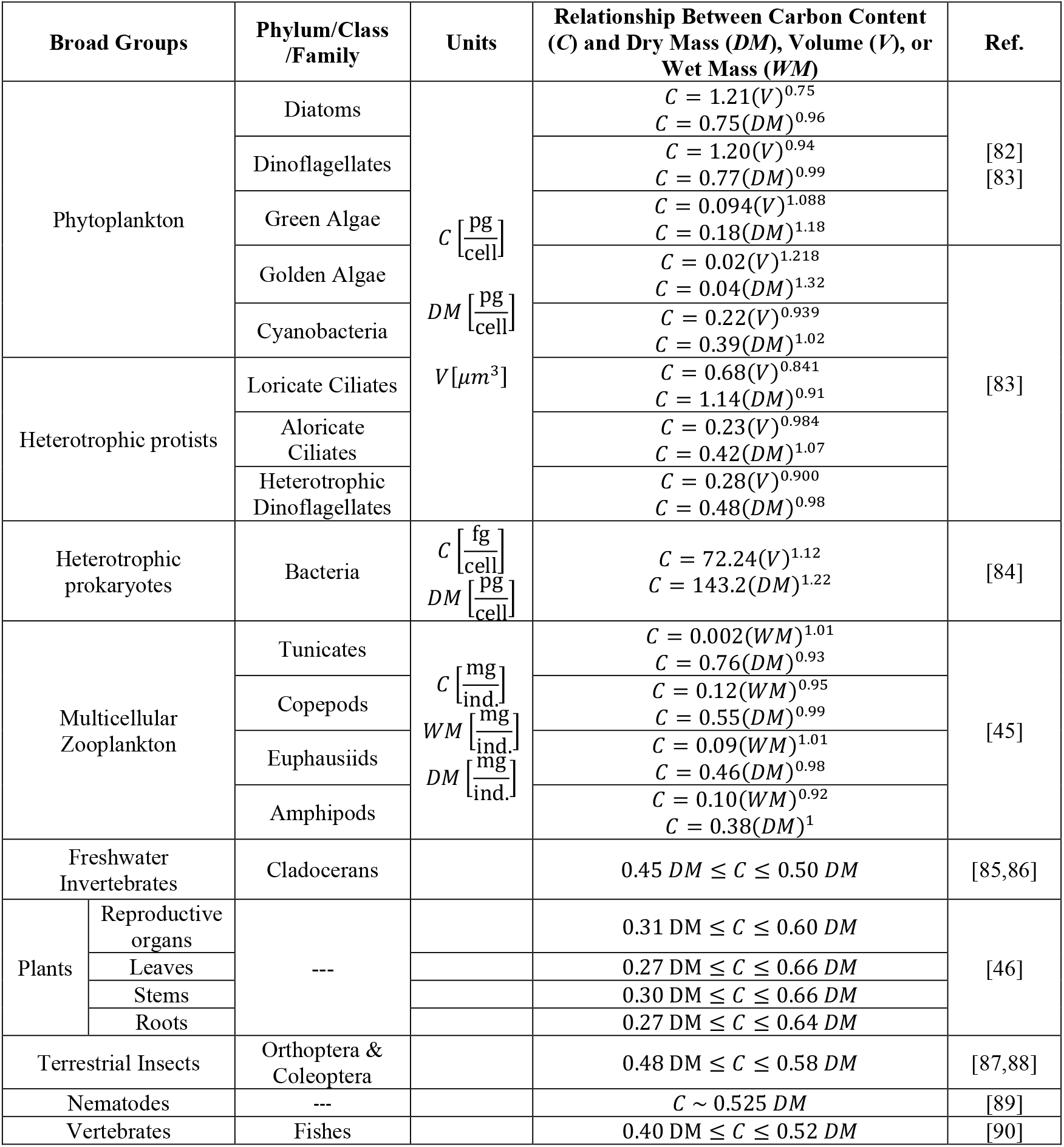
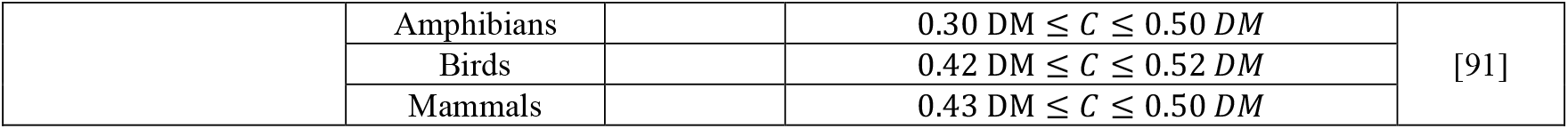
Mass scaling of carbon content across the kingdoms of life. For details on how these equations were derived, see the section “Standardization of Respiration Rates and Carbon Content Across Units and Temperature” in the Supplementary Section II.4.

**Table S4:**
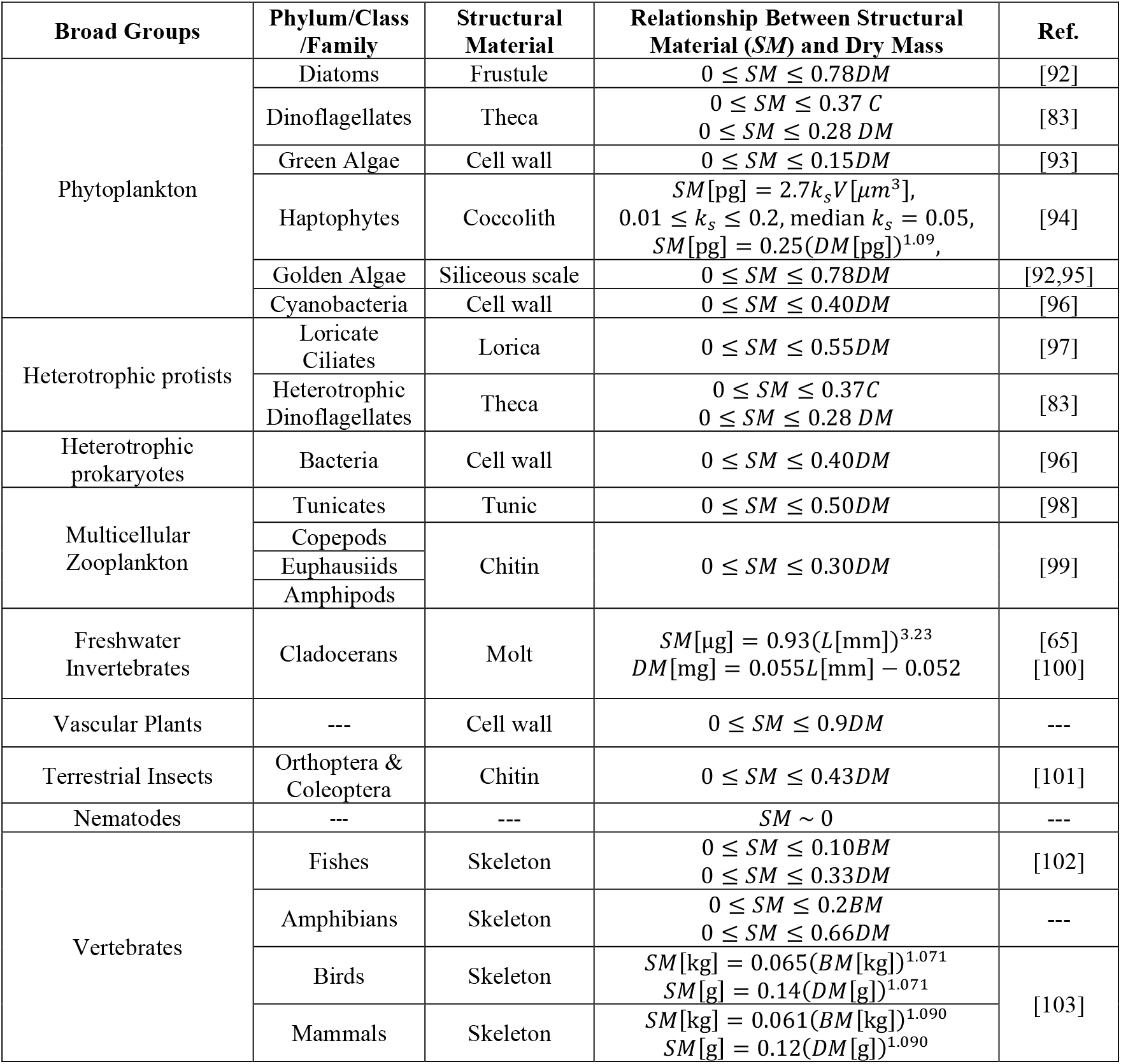
Structural material-to-dry mass ratios across the kingdoms of life. When structural material mass is described as ranging from zero to a specific value, it indicates that the structural material cannot be definitively excluded, as it may be metabolically active. Therefore, a range of values starting from zero to a maximum is considered in the numerical simulation of the carbon-to-ATP ratio.

**Table S5:**
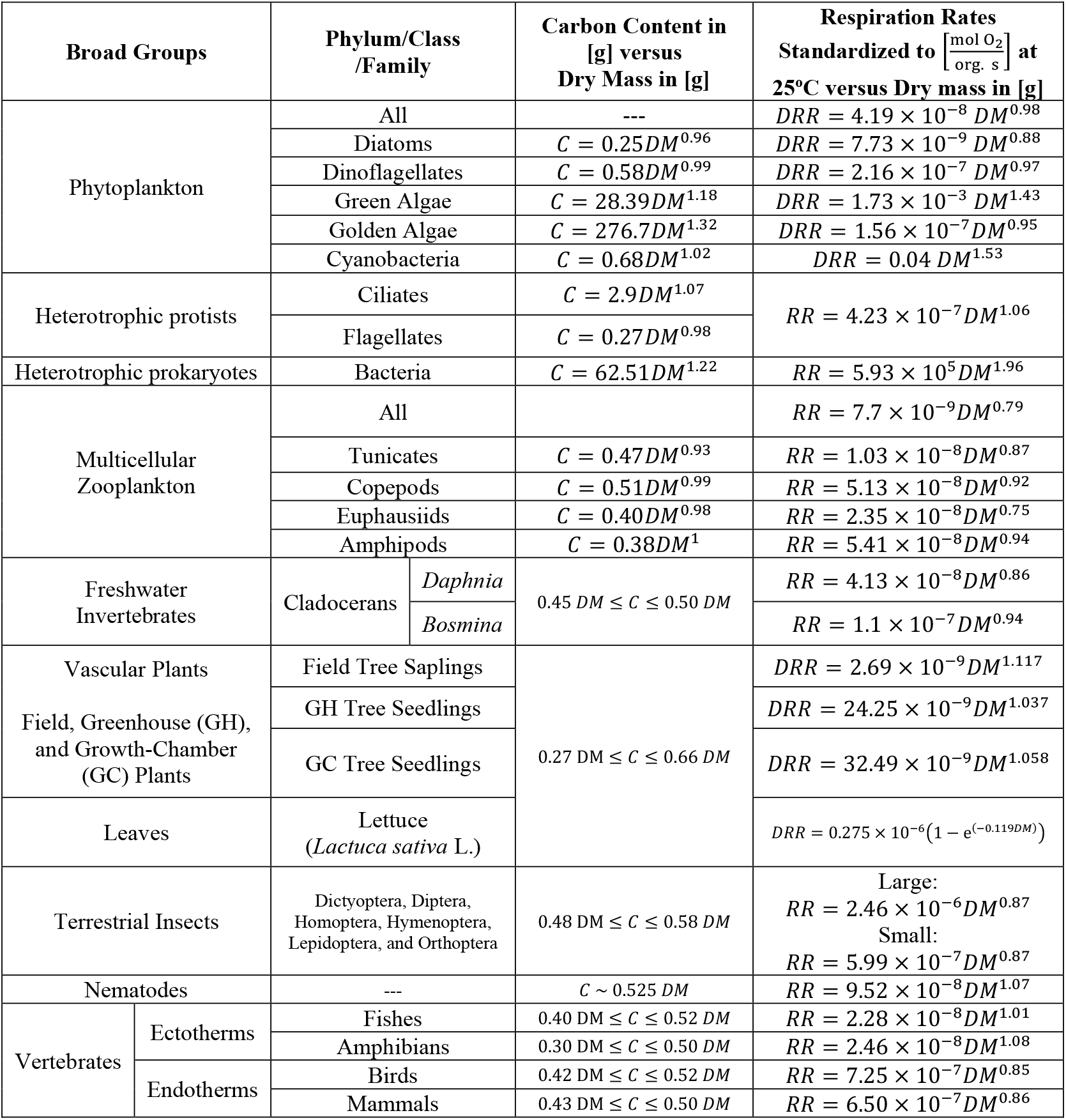
Respiration rates standardized 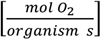 at 25°C, with carbon contents and dry mass expressed in grams.

#### II.4. Standardization of Respiration Rates and Carbon Content Across Units and Temperature

To enable direct comparison across datasets, respiration rates (Table S2) and carbon content (Table S3) were standardized to the same units and normalized to the reference temperature provided in Table S5; here we elaborate how this standardization was mediated.

##### II.4.1. Respiration Rates

###### Diatoms and Cyanobacteria

The data are taken from [13], where dark respiration rates (DRR) are reported in units of watts per kilogram of wet mass, and body mass (BM) is given in picograms at 25 °C. We first converted DRRs to watts per organism by multiplying the original unit by BM. A reduced major axis (RMA) regression was then applied to obtain the equation presented in Table S2. The standardized equations in Table S5 are expressed in mol O_2_ per organism per second at 25 °C, with organism mass reported as dry mass (DM). For this conversion, we assumed that 1 ml O_2_ = 20 J, the molar volume of O_2_ = 22,400 ml/mol at STP, and DM = 0.3 BM.

###### Dinoflagellates, Green Algae, and Golden Algae

The data are taken from [72], where DRRs are reported in pl O_2_ per organism per hour, and organismal carbon content is given in picograms. An RMA regression was applied to derive the equation presented in Table S2. Although DRR versus carbon content is a more direct metric for estimating C:ATP ratios, we converted carbon to DM in order to provide equations consistent with those of other organisms for comparative purposes. For this conversion, we used the relationship *C*[pg]≅1.59×(*DM*[pg])^0.96^ developed for phytoplankton in [82]. The resulting equations are reported in Table S2. Finally, DRRs and DM were standardized to the units used in Table S5, applying the conversion factors described in the previous paragraph.

###### Heterotrophic Protists and Bacteria

The equations are taken from [18,73], where RRs are reported in watts per organism at 20 °C, with BM expressed in grams. To standardize the units, we applied the following conversions: 1 ml O_2_ = 20 J, the molar volume of O_2_ = 22,400 ml/mol at STP, DM = 0.3 BM, and Q_10_ = 2.5.

###### Multicellular Zooplankton

The data are taken from [14], where RRs are reported in microliters of O_2_ per organism per hour at 15 °C, and carbon content is given in micrograms. We applied an RMA regression to derive the relationship between RR and carbon content. For consistency with other organisms, we converted carbon content to DM using the zooplankton-specific equations provided in [45]. Finally, we standardized the units to those used in Table S5 and adjusted the temperature to 25 °C using Q_10_ = 2.5.

###### Freshwater *Daphnia* and *Bosmina*

The equations are taken from [65], where average RRs are reported for *Daphnia* species and for *Bosmina longirostris* in microliters of O_2_ per organism per day at 20 °C, expressed as a function of DM in micrograms. The units were standardized to those used in Table S5 using the conversion factors described in the previous paragraphs.

###### Vascular Plants

The equations are taken from [75], where whole-plant RRs of various plant types are reported in nmol O_2_ per plant per second at 24 °C as a function of DM in grams. In Table S5, these equations are converted to mol O_2_ per second and standardized to 25 °C.

###### Leaves

The equation is taken from [67], where RR for lettuce (*Lactuca sativa L*.) is reported in micromoles O_2_ per plant per second at 25 °C as a function of DM in grams. For consistency, RR was converted to moles O_2_ per second and presented in Table S5.

###### Insects

The equations are taken from [68], where RRs during flight are reported in mm^3^ O_2_ per organism per hour at 22 °C as a function of BM in milligrams. To convert mm^3^ O_2_ to moles O_2_, as reported in Table S5, it was assumed that 1 mole O_2_ = 22.4 × 10^6^ mm^3^ at STP. All other units were standardized to those in Table S5 using the same assumptions described previously.

###### Nematodes

The data are taken from [69], considering only species from oxic sites with respiration measured under oxic incubation. RRs were reported in nmol O_2_ per organism per day at 10 °C as a function of BM in micrograms. BM was first converted to DM using the relation DM = 0.3 BM, and an RMA regression was then applied to derive the equation presented in Table S2. Finally, units were standardized to those reported in Table S5 using the conversion factors described earlier.

###### Vertebrates

The data are taken from [70], where maximal RRs are reported in ml O_2_ per organism per hour at various temperatures as a function of BM in grams. Temperatures were normalized by converting ectotherm values to 25 °C and endotherm values to 38 °C using Q_10_ = 2.5. An RMA regression was then applied to obtain the equations presented in Table S2. Finally, units were standardized to those in Table S5 using the conversion factors described earlier.

##### II.4.2. Tissue-specific Respiration Rates in Animals

In Figure 1 of the paper, we present an example of modeling tissue-specific C:ATP ratios for rat brain, heart, kidney, and liver. To estimate tissue-specific respiration rates, we first calculate the whole-body respiration rate using the mammalian scaling equation provided in Table S5. We then obtain organ-specific fractions of total oxygen consumption in rats from the literature [49]. Tissue-specific respiration rates are calculated by multiplying the whole-body respiration rate by the corresponding organ-specific fraction (e.g., 3% for brain). The ranges of whole-body dry mass used to estimate whole-body respiration rates, as well as the ranges of organ-specific dry mass used to estimate tissue carbon content in rats, are provided in Supplementary Excel File 1.

##### II.4.3. Carbon Contents

###### Diatoms, Dinoflagellates, and Green Algae

The equations are taken from [82], where carbon content per organism (pg) is reported as a function of cell volume (μm^3^). Dry mass (pg) is also reported as a function of cell volume (μm^3^). Combining these two equations allowed us to establish the relationship between carbon content and dry mass, as presented in Table S3. Both variables were then converted to grams to match the units reported in Table S5.

###### Golden Algae, Cyanobacteria, Ciliates, and Heterotrophic Dinoflagellates

The equations are taken from [83], where carbon content (pg cell^−1^) per organism is reported as a function of cell volume (µm^3^), as shown in Table S3. We first converted volume to dry mass (DM) using the equation 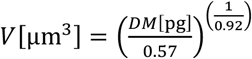, developed in [1] for unicellular organisms, with the resulting equations provided in Table S3. We then standardized the units to match those in Table S5, where both carbon content and DM are expressed in grams.

###### Heterotrophic Prokaryotes

The equation is taken from [84], where carbon content in femtograms per organism is reported as a function of cell volume in μm^3^ (see Table S3). We first converted volume to DM using the relation 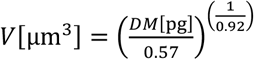, as provided in Table S3. We then standardized both carbon content and DM to grams, consistent with the units reported in Table S5.

###### Multicellular Zooplankton

The equations are taken from [45], where carbon content per organism (mg) is reported as a function of wet mass (mg). Separately, dry mass (mg) is reported as a function of wet mass (mg). By combining these two equations, we derived the relationship between carbon content and dry mass, as shown in Table S3. Both carbon content and dry mass were then standardized to grams, consistent with the units reported in Table S5.

### III. Proton Leak in Mitochondria

Not all oxygen molecules respired are necessarily converted into ATP [104]. Unregulated basal proton leak (PL) through adenine nucleotide translocases (ANT) [104], and water wires (WW) [22], as well as regulated inducible PL through uncoupling proteins (UCPs) and ANTs [22,104], dissipates a significant amount of energy as heat without ATP production. Presence of UCPs have been reported in mammals, birds, reptiles, amphibians, fishes, insects, plants, fungi, and protozoa [105], and PL has also been observed in bacterial cells [106,107]. PL increases nonlinearly with inner mitochondrial membrane potential [21,108] and correlates positively with the standard metabolic rate of organisms [108]. Due to the limited data on the relationship between mitochondrial PL and metabolic rate across all kingdoms of life, we are not using scaling equations to estimate PL at this time. This is primarily because the electrochemical potential of the inner mitochondrial membrane (IMM) is highly dynamic, fluctuating in time and space under both physiological and stress conditions [109], as well as during the fission and fusion of mitochondrial networks [110–113]. For example, the potential of the IMM is not homogeneous; it is higher in the cristae relative to the inner boundary membrane [114] and is elevated at the sites of ATP synthase molecules due to the intrinsic molecular electrostatic potential of this enzyme [115]. Additionally, the membrane potential is dynamically regulated through various chemical pathways and electrical feedback mechanisms [116], making it challenging to develop an equation that accurately represents the relationship between respiration rate and proton leak. As such, we rely on percentage estimates reported in the literature, considering basal PL to account for at least 20% to 50% of the standard metabolic rate [21,22].

### IV. Stoichiometric Ratios of ATP/O_2_ in Oxidative Phosphorylation

The maximum number of ATP molecules produced per oxygen atom consumed 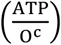 in oxidative phosphorylation and glycolysis pathways [10,16] is determined by the equation:

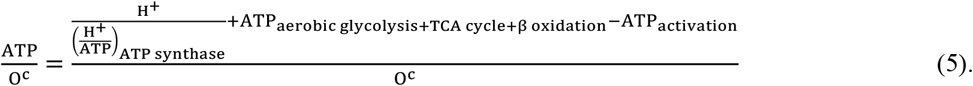

Here, the 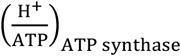 ratio represents the number of protons translocated through ATP synthase to produce one ATP molecule. All other parameters depend on the specific metabolic pathway. For instance, during the oxidation of palmitate: O^c^ = 46, total H^+^ = 392, ATP_activation_ = 2, and ATP_TCA cycle+β oxidationation_ = 8 [10,16]. The 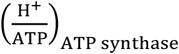 ratio depends on the number of c-subunits in the ATP synthase complex. For example, the ATP synthase in the mitochondria of *Saccharomyces cerevisiae* [117], *Yarrowia lipolytica* [118], and *Euglena gracilis* [119] contains 10 c-subunits, while in human [120], bovine [121], and pig [122] mitochondria, it contains 8 c-subunits [123,124]. Due to the absence of data on the number of c-subunits in the mitochondrial ATP synthase of all the organisms studied in this paper, we assume that mitochondrial ATP synthase in unicellular organisms contain 10 c-subunits, while mitochondrial ATP synthase in multicellular organisms contain 8 c-subunits. In heterotrophic bacteria, the number of c-subunits ranges from 9 to 17 [123,124].

For each ATP produced in mitochondria, one inorganic phosphate (Pi) is transported into the mitochondrial matrix via phosphate carriers (PiC), accompanied by one proton [125]. Consequently, we must add three additional protons to the total from each c-subunit [16]. This results in 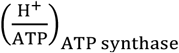 ratios ranging from approximately (8+3)/3 ∼ 3.66 and (10+3)/3 ∼ 4.33. In bacterial ATP synthase, the F_1_ portion of the enzyme is located in the cytoplasm [126], where inorganic phosphate is already present. Therefore, no phosphate carrier (PiC) is required. As a result, the proton-to-ATP ratio for ATP synthase 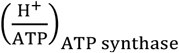 ranges from 9/3 = 3 to 17/3 ∼ 5.66, depending on the number of c-subunits in the ATP synthase. Assuming that the metabolic pathways involved in oxidative phosphorylation and glycolysis are largely conserved across all kingdoms of life, with differences primarily in the number of c-subunits in ATP synthase, the Table S7 presents the minimum and maximum values of 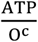 for c-subunits ranging from 8 to 17. To simplify future calculations, we double the values to obtain the ATP per molecule of oxygen 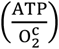 in Table S6.

**Table S6:** Minimum and maximum ATP yield per oxygen molecule consumed in oxidative phosphorylation and aerobic glycolysis, associated with the number of c-subunits in ATP synthase and the metabolic pathway.

| Number of c-subunits | Considering $H^+$ associated with PiC? | Minimum $\frac{ATP_{yield}}{O_{2-c-max}} \rightarrow$<br>succinate to malate pathway | Maximum $\frac{ATP_{yield}}{O_{2-c-max}} \rightarrow$<br>glycogen to $CO_2$ pathway |
| --- | --- | --- | --- |
| 8 | Yes | 3.27 | 5.74 |
| 9 | No | 4 | 6.83 |
| 10 | Yes | 2.77 | 4.99 |
|  | No | 3.6 | 6.23 |
| 11 | No | 3.27 | 5.74 |
| 12 |  | 3 | 5.33 |
| 13 |  | 2.77 | 4.99 |
| 14 |  | 2.57 | 4.7 |
| 15 |  | 2.4 | 4.43 |
| 16 |  | 2.25 | 4.21 |
| 17 |  | 2.12 | 4.01 |

### V. Photosynthesis Rate, Proton Leak, and 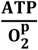 Ratio in Chloroplast

In photoautotrophic organisms, the ATP synthase in chloroplasts contains 14 c-subunits, as reported for *Spinacia oleracea* [127], *Pisum sativum* [128], and *Triticum aestivum* [129], whereas in *Synechococcus* cyanobacteria, it ranges from 13 to 15 c-subunits [123,124]. Although no crystallographic or microscopic studies have reported an ATP synthase in chloroplasts with fewer than 14 c-subunits, it has been suggested that the chloroplast ATP synthase in *C. reinhardtii* may have 13 c-subunits instead of the usual 14. This suggestion is based on observed differences in the migration pattern of proteins in SDS-PAGE gels [130].

In chloroplasts, during non-cyclic photophosphorylation or linear electron flow (LEF) under standard conditions, twelve protons are pumped into the thylakoid lumen for each oxygen molecule produced during photosynthesis [131], yielding an 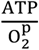 ratio of approximately 12 ÷ ((14 or 13) ÷ 3) ∼ 2.57 to 2.77. However, as discussed earlier and previously reported for mitochondria [10,16], each inorganic phosphate (Pi) used in ATP synthesis is accompanied by the transport of one proton from the cytoplasm into the chloroplast via the phosphate carrier (PiC), a factor often overlooked in chloroplast studies. Accounting for this, the adjusted 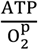 ratio ranges approximately from 2.12 to 2.25. Under standard conditions, chloroplasts require at least three ATP molecules per two NADPH molecules to sustain carbon fixation alone [17]. The additional ATP necessary for other processes is generated through cyclic photophosphorylation or cyclic electron flow (CEF), which does not involve oxygen and supports the higher ATP/NADPH ratio needed for the Calvin cycle [132], CO_2_-concentrating mechanisms [133–135], and other intra-chloroplast processes. Determining the proportion of ATP produced via CEF versus LEF depends on the rates of both pathways. Literature reports that the CEF-to-LEF ratio can range from 3% [132] to 14% [131] and up to 60% [27]. However, it has been reported that the rate of electron transfer per second per PSI complex in CEF can reach up to 210 e^−^ s^−1^ PSI^−1^, suggesting that CEF can potentially operate as fast as LEF [136–138]. Additionally, the pseudo-cyclic pathway, where molecular oxygen acts as the terminal electron acceptor in the electron transport chain, can generate ATP at rates as high as 20% of LEF [27]. Thus, it is reasonable to assume that 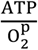 ranges from 2.12 + 2.12 + 0.2(2.12) = 4.66 to 2.25 + 2.25 + 0.2(2.25) = 4.95, accounting for the combined contributions of all LEF, CEF, and pseudo-CEF pathways.

Proton leakage across the thylakoid membrane of chloroplasts has been observed under specific conditions, such as low inorganic phosphate levels in isolated spinach and lettuce thylakoids [139,140]. However, the extent of this leakage remains poorly studied, and the available literature reports inconsistent values. For example, in *Spinacia oleracea*, uncoupling of protons via the thiol reagent dithiothreitol or light has been reported to dissipate 20–30% of ATP yield [141]. In contrast, a study on *Arabidopsis* leaves found the ion leakage to be negligible [142]. Additionally, in young spinach leaves, ion leakage (though unspecified as to the type of ions) increased under dark conditions when the electrochemical proton gradient exceeded 190 mV [143]. Therefore, our model assumes that proton leak dissipates between 0% and 30% of the ATP production rate in the chloroplast.

To estimate the total rate of ATP production in photoautotrophic organisms, it is necessary to account for all ATP-producing pathways, including mitochondrial pathways, glycolysis, and chloroplast-mediated pathways. A key missing element is the relationship between respiration and photosynthesis rates in microalgae and higher plants. The ratio of dark respiration rate to light-saturated photosynthesis rate 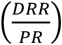, defined as 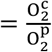, varies across species. For actively growing cells, this ratio ranges from 0.08 to 0.6 in Dinophyceae (mean ∼ 0.35±0.17), 0.013 to 0.50 in Bacillariophyceae (mean ∼ 0.16±0.06), 0.043 to 0.63 in Chlorophyceae (mean ∼ 0.13±0.07), 0.03 to 0.64 in Prymnesiophyceae (mean ∼ 0.16±0.04), and 0.004 to 0.17 in Cyanophyceae (mean ∼ 0.10±0.07) [15,28,29]. It is observed that this ratio is independent of cell size [15] and light intensity [23,24]. For vascular plants, we use ratios associated with Chlorophyceae.

### VI. Steady-State ATP Turnover Time

As discussed in the main text, the steady-state ATP turnover time depends on the diffusion time of ATP from its site of production to its site of consumption. Here we explore quantitatively the factors known to affect ATP delivery time.

#### VI.1. ATP Diffusion Coefficient

ATP molecules are produced in the mitochondrial matrix, where the viscosity is twice that of pure water, and their diffusion is significantly hindered by the presence of cristae membranes that vary across different respiratory states [144]. Subsequently, ATP must traverse the highly selective mitochondrial membrane, where the ATP-ADP exchange rate depends on the membrane’s electrochemical potential [145]. At steady state, in liver mitochondria depolarized to different membrane voltages, the ATP exchange rate ranges from near zero at −100 mV to 0.5 mM ATP per minute per milligram of protein at −145 mV [145]. Various studies have reported a range of diffusion coefficients for ATP in the cytoplasm. Effective diffusion coefficients as low as 100 μm^2^/s [146] have been observed when accounting for path length and interactions with cytoplasmic components. Other reported values include 200 μm^2^/s [147], 236 μm^2^/s [148], and a temperature- and pH-dependent diffusion coefficient ranging from 153 μm^2^/s at pH 7.31 and 5°C to 480 μm^2^/s at pH 7.18 and 40°C [149]. Furthermore, the diffusion coefficient is time-dependent; over long timescales, the translational diffusion of a molecule the size of ATP can be reduced to as little as 10–50% of its value in dilute solution [150]. The average cytoplasmic microviscosity reported for HeLa cells is as high as 86 cP [151], nearly two orders of magnitude greater than the viscosity of water at 20 °C. According to the Stokes–Einstein relationship in the low Reynolds number regime ( 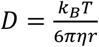, where *D* is the diffusion coefficient, *k*_B_ is the Boltzmann constant, *η* is the viscosity, and *r* is the radius of the diffusing spherical particle), such high cytoplasmic microviscosity results in a diffusion coefficient that is two orders of magnitude lower than in water. Experimentally tracing all ATP molecules from their site of production to their site of consumption - anywhere within the cell - is not feasible. However, the above gathered evidence indicates that the ATP diffusion coefficient depends on multiple factors, including distance, time, microviscosity, molecular crowding, membrane exchange rates, pH, temperature, cell type, and more. These dependencies suggest that the diffusion coefficient can vary widely and may, under certain conditions, be significantly lower than commonly reported values. However, for our purposes, we adopt a value of 100 μm^2^/s.

#### VI.2. Average ATP Diffusion Time

We investigate two modes of ATP transport: (1) diffusion, occurring in both prokaryotic and eukaryotic cells, and (2) mitochondrial motility, where ATP is delivered directly by mitochondria to various cellular compartments. In the case of free diffusion, the average diffusion time in three-dimensional space is estimated by 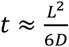, where *L* is the diffusion distance (approximated here as the cell length) and *D* is the diffusion coefficient. For cells with typical sizes - 1 μm for bacteria, 100 μm for ciliates [43], and 22 μm for adult human cells [36] - the corresponding ATP diffusion times are approximately 0.002 s, 17 s, and 0.8 s, respectively. However, in the cytoplasm, ATP transport does not occur via free diffusion. Instead, it follows a drift-diffusion behavior, influenced by ATP concentration gradients across various organelles and compartments. ATP is actively targeted to specific sites based on metabolic demand. The average time required for ATP molecules to reach their target in 3D space under such conditions can be approximated by 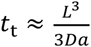, where *a* is the radius of the target region, assuming *L* ≫ *a* [152]. Taking *a* = 0.1*L*, the estimated ATP delivery times are approximately 0.03 s for bacteria, 330 s for ciliates, and 16 s for human cells. These back-of-the-envelope calculations illustrate the two extremes of ATP diffusion time across differently sized organisms, influencing the average ATP turnover time. In the second scenario, ATP is delivered by mitochondrial motility. Mitochondria are motile organelles with experimentally observed speeds ranging from 24.2 nm/s in dendrites to 185 nm/s in axons [153]. Based on these velocities, ATP delivery times via mitochondria would be roughly 540–4132 s for a typical ciliate cell and 119–909 s for a typical adult human cell. This clearly shows that mitochondrial motility is not an efficient mechanism for rapid ATP delivery and that repositioning mitochondria within the cell is time-consuming. Taken together, this evidence supports the existence of ATP concentration gradients as a function of distance from mitochondria, varying with physiological conditions - a phenomenon well documented in the literature [146,154–157].

In animals, ATP levels as low as 1.18 × 10^−4^ g ATP / g dry mass have been reported in amphipods in saltwater at 9°C [158], whereas values as high as 4.46 × 10^−1^ g ATP / g dry mass have been observed in the brain of nondiapausing pupae of *H. armigera* [159], illustrating a difference of more than three orders of magnitude. One of the main reasons for these differences is the mechanism by which oxygen is delivered to cells via closed, open, or mixed circulatory systems [160]. In general, oxygen tension decreases with increasing distance from the vessel wall, as observed in various tissues across different organisms - for example, in the arterioles of the rat mesentery [161]. A further reduction in oxygen availability can occur during intense activity; for example, in Weddell seals during diving, blood flow to organs such as the pancreas, liver, and heart decreases markedly [162], leading to reduced oxygen supply and consequently lower ATP levels. Other potential contributors to these differences include the broad range of cell sizes in animals and their varied metabolic activities, as discussed in the main text.

Both experimental and theoretical evidence suggest that oxygen molecules diffuse more rapidly in lipids than in cytosolic and interstitial fluids, due to their higher solubility in lipids, which helps overcome diffusion barriers over long distances [163]. This suggests that in animal cells - similar to heterotrophic protists - some mitochondria should localize near the cell membrane, while others position themselves close to energy-demanding compartments such as the nucleus [164]. Animals with high respiration rates, such as insects, tend to have more mitochondria than those with lower respiration rates, such as fish. This abundance enables greater flexibility in mitochondrial spatial distribution, allowing for more efficient positioning across the cell, thereby minimizing ATP delivery time and, consequently, ATP turnover time.

### VII. On the Size-Dependence of C:ATP

As shown in the Supplementary Section I, based on the compiled C:ATP data, there is a statistically significant relationship between organism or tissue dry mass and the C:ATP ratio for several of the organisms and tissues examined, although some exhibit positive correlations while others show negative correlations. As discussed in the main text, we cannot fully elaborate on these patterns at this stage because we do not yet have an explicit model for size-dependent ATP turnover time. However, if we assume a constant ATP turnover time, we can plot our theoretically modeled C:ATP versus dry mass for the organisms studied here, where the positive or negative relationships arise simply from the scaling exponent of carbon content versus dry mass divided by the scaling exponent of respiration rate versus dry mass (Table S5). The result of this analysis is shown in Figure S6, which also illustrates how much variation in C:ATP is attributable to size differences across the observed size range for each organism.

**Figure S6:**
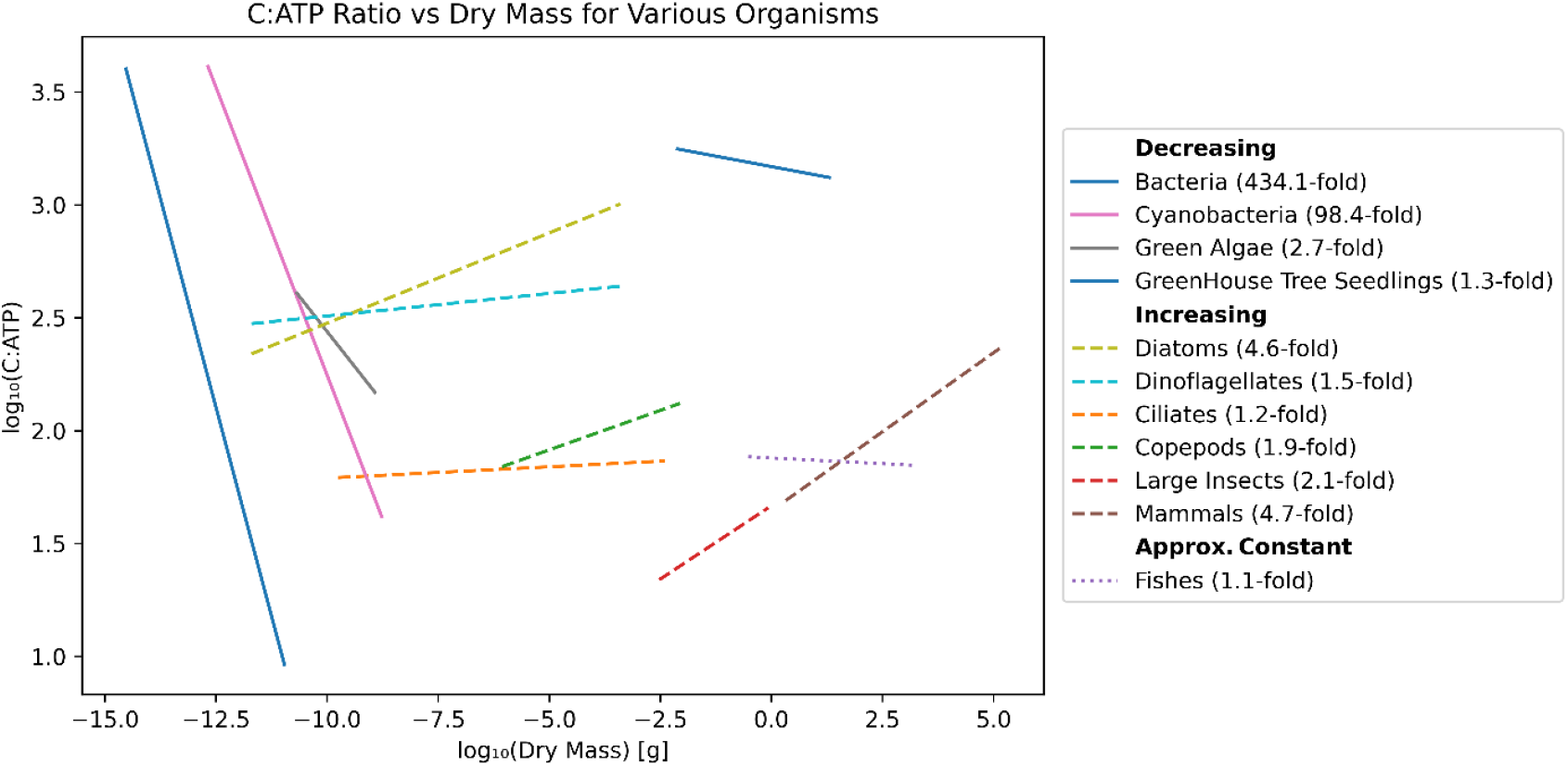
This figure shows the relationship between log(C:ATP) and log(Dry Mass [DM]) based on our model. In this analysis, only carbon content and respiration rate are modeled as functions of dry mass; all other parameters are fixed at their median values. The results indicate that in some organisms, C:ATP increases with DM, whereas in others it decreases or remains nearly constant. The legend reports the fold change in C:ATP across the full DM range, from smallest to largest sizes. Notably, bacteria and cyanobacteria show large variations because the exponent for respiration rate versus DM in these groups is much higher than that for carbon versus DM.

We also performed a sensitivity analysis on the parameters in Eqs. (2) and (3) while keeping ATP turnover time constant. For each organism, we fixed all parameters at their baseline values. We then varied one parameter at a time across its prescribed plausible range (sampled on a uniform grid of 100 values) and recomputed the model-predicted C:ATP for each value. Sensitivity was summarized as a fold-change metric for each parameter, defined as “max(C:ATP)/min(C:ATP)” over that range. Figure S7 presents a two-dimensional heat map: one dimension shows variations in C:ATP caused by parameter changes within groups (row-wise comparison, blue map), and the other shows variations across groups (column-wise comparison, orange map). For example, to assess the importance of parameters influencing C:ATP in fish, one should examine the blue colors in the fish row. This indicates that 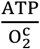 has the strongest effect (darkest blue), whereas dry mass (DM) has the weakest effect (lightest blue). Likewise, to compare the effect of DM on C:ATP across groups, one should examine the orange colors in the DM column. This column shows that bacteria (darkest orange), followed by cyanobacteria, are the most strongly influenced by DM among the organisms.

**Figure S7:**
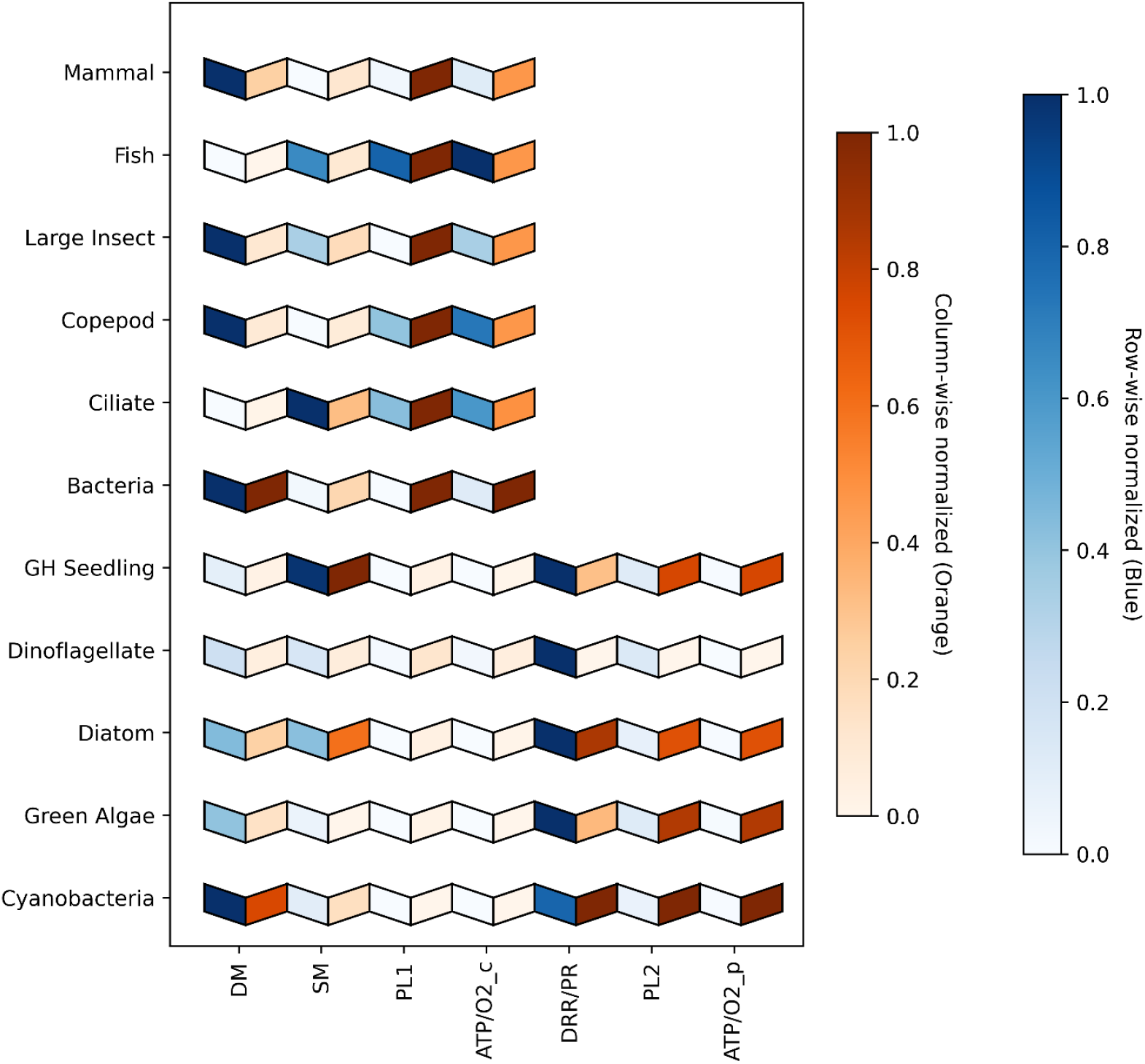
Two-dimensional sensitivity analysis of parameters in Eqs. (2) and (3) under the assumption of constant ATP turnover time. The heat map displays (i) within-group variation in C:ATP in response to parameter changes (row-wise, blue scale) and (ii) across-group variation for each parameter (column-wise, orange scale).

